# Nested small open reading frames are translated from alternative transcripts

**DOI:** 10.1101/2024.10.22.619581

**Authors:** Haomiao Su, Samuel G. Katz, Sarah A. Slavoff

## Abstract

Overlapping genes were thought to be essentially absent from the human genome until the discovery of hundreds of translated, frameshifted internal open reading frames (iORFs) within annotated protein coding sequences (CDS). This would suggest that some human genes encode two completely different proteins; however, it is unclear how iORF-encoded proteins are translated, and whether they are broadly functional. We demonstrate that “non-coding” alternative transcripts lacking a complete protein coding sequence (CDS) are required for iORF translation. We also demonstrate that iORFs such as the conserved and antiapoptotic *DEDD2* iORF encode proteins with functions distinct from the annotated proteins they overlap. This work thus provides a molecular and functional basis for dual coding of overlapping ORFs in human genes.

## Introduction

Small open reading frames (smORFs) encoding microproteins of fewer than 100 amino acids were previously excluded from the human genome annotation. However, thousands of human smORFs are now known to be translated(*1, 2*), and hundreds are essential for cell proliferation or survival (*3, 4*). Furthermore, smORFs have been implicated in diseases including cancer, metabolic disorders and aging(*4-6*). Despite this progress, the extent to which smORF translation results in production of functional microproteins remains under debate. In particular, hundreds of internal, frameshifted smORFs (iORFs) have been mapped deep within protein coding sequences (CDSs) in mRNA, but their translation is expected to be repressed by recognition of the CDS start codon by ribosomes. Dual translation of a handful of iORFs and the overlapping CDS has been reported(*7, 8*), but this process – termed leaky scanning (*9*) – is only possible in specific contexts. Presenting an even greater conceptual hurdle, since iORFs are frameshifted relative to the CDS that they overlap, their amino acid sequences are completely different than the canonical proteins that they overlap. The existence of iORFs would therefore imply that some human genes encode two different proteins - a phenomenon known to occur in viruses, but not eukaryotes – and human iORF translation cannot be broadly rationalized. We therefore sought to establish a mechanistic basis for human iORF expression and to expand evidence for functional microproteins encoded in iORFs.

We propose that some iORFs have been mis-mapped within protein CDSs, and instead are translated from alternative transcripts that lack the upstream CDS start codon, some of which have not previously been detected (Fig. 1A). Supporting this idea, a majority of human genes exhibit non-coding alternative transcripts that lack the complete annotated CDS (*10, 11*). We suggest that a subset of these transcripts may not be non-coding at all, but instead encode iORFs and thus change the protein product of the gene. Expression of two different proteins from the same gene via alternative transcripts has previously been reported: for example, transcripts arising from alternative transcription start sites (TSS) in the *CDKN2A* gene encode INK4A and ARF, which share no amino acid sequence (*12, 13*). Alternative transcripts have also been invoked to explain iORF expression: we previously reported that the human *DEDD2* gene generates alternative transcripts that encode either the annotated, nuclear DEDD2 protein or a mitochondrial microprotein (*14*). In this work, we provide evidence that translation of iORFs in human cells often requires alternative transcripts, and further characterize the human *DEDD2* gene, which encodes both the pro-apoptotic canonical DEDD2 protein and an antiapoptotic iORF-encoded microprotein in alternatively spliced transcripts.

**Fig 1.**
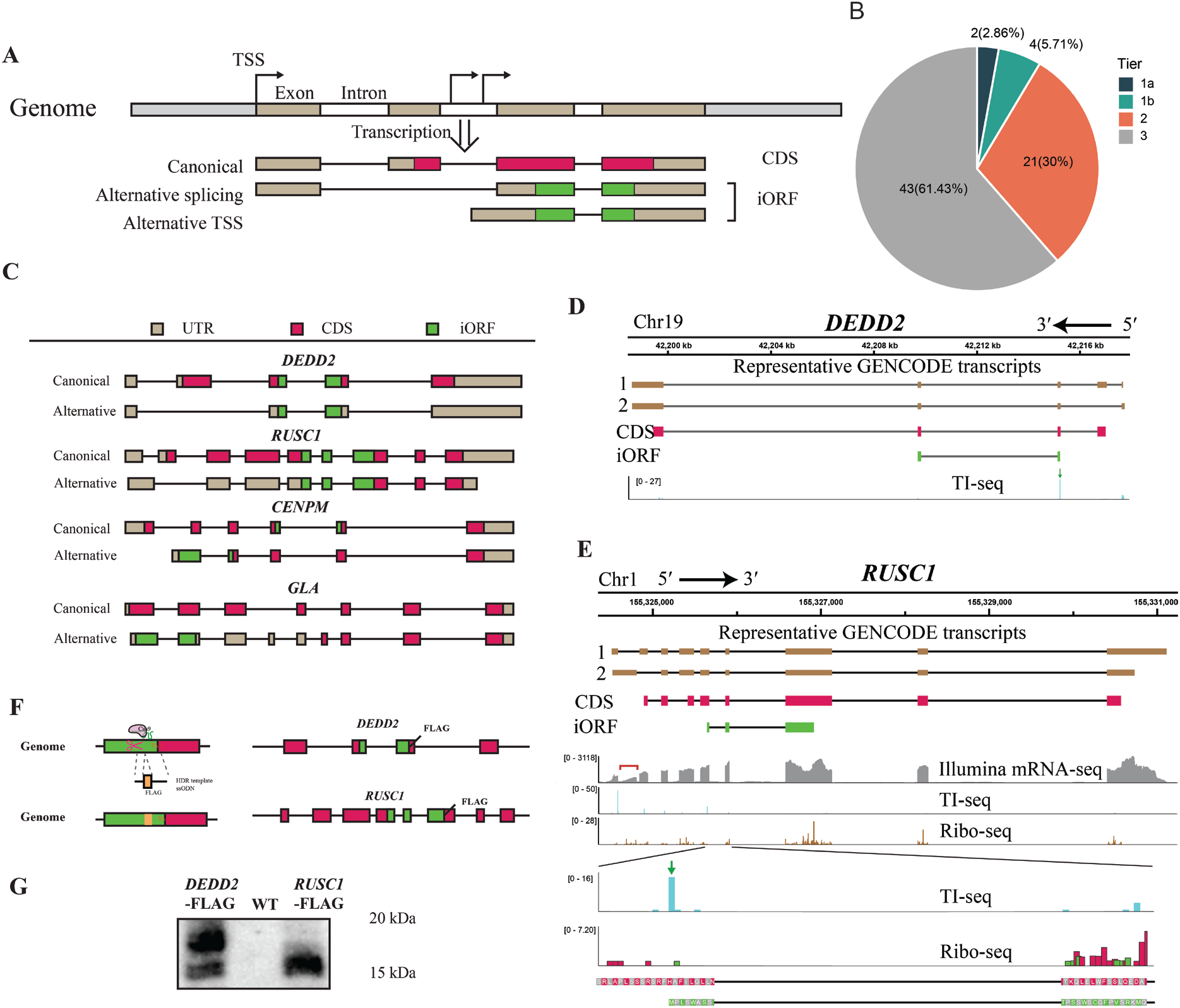
Internal, overlapping ORF (iORF) expression from alternative transcripts. (**A**) iORFs that cannot be expressed via specialized translation downstream of an annotated start codon may instead originate from alternative transcripts that lack the upstream start. (**B**) Pie chart illustrating number of iORF target genes in each confidence tier (1a, 1b, 2, 3) subjected to ORF capture-seq in this work. (**C**) Schematic of iORFs and CDSs in GENCODE canonical (upper panel) and alternative transcripts (lower panel) for four representative genes examined in this study. (**D**) Alternative transcript encoding an iORF in *DEDD2*. Tracks show the genomic locus above, canonical (1) and alternative (2) transcripts, coding regions for the annotated RUSC1 coding sequence CDS (magenta) and iORF (green), and translation initiation site sequencing (TI-seq) signal (cyan). (**E**) Alternative transcript encoding an iORF in *RUSC1*. Tracks show the genomic locus above, canonical (1) and alternative (2) transcripts, coding regions for the RUSC1 CDS (magenta) and iORF (green), mRNA-seq reads (gray), TI-seq signal (cyan), and Ribo-seq. The red bracket highlights RNA-seq reads specific to the alternative transcript. The green arrow points to the iORF start codon in the TI-seq data. Ribo-seq reveals reads specific to the iORF reading frame (green) distinct from the CDS reading frame (magenta). (**F**) Strategy for CRISPR/Cas9 homology directed repair genomic knock-in (KI) of HA-FLAG epitope tags at the 3’ ends of the *DEDD2* and *RUSC1* iORFs to report on endogenous iORF expression. (**G**) Anti-FLAG Western blot of immunoprecipitates from clonally selected DEDD2-FLAG and RUSC1-FLAG KI cells versus parental HEK293T (WT).

## Results

### Evidence for iORF translation from annotated alternative transcripts

In order to assess the translation mechanism of iORFs in human cells, we first required a list of high-confidence translated iORFs. While iORF translation has been previously reported using both ribosome profiling and mass spectrometry, and gold-standard translated smORF catalogs are available(*15*), it is likely that iORFs are underrepresented in these resources. This is because proteomics under-detects *bona fide* smORF-encoded microproteins(*16*), and ribosome profiling data analysis prioritizes three-nucleotide periodicity in a single reading frame, which can obscure dual translation of iORFs(*17*). We therefore integrated and reanalyzed data from multiple resources to identify translated iORFs with varying levels of evidence, which we assigned to tiers (Fig. 1B). Tier 1a iORFs are the highest confidence and are supported by ribosome profiling (RIBO-seq) evidence in a gold-standard human translatome resource (*15*) as well as proteomic evidence (two unique peptide-spectral matches identified in human leukocyte antigen I (HLA-I) immunopeptidomics data available in PeptideAtlas) (*18*). Tier 1b iORFs are supported by RIBO-seq and at least one unique peptide. Tier 2 iORFs are supported only by RIBO-seq evidence, and Tier 3 iORFs are not identified in either resource. We included Tier 3 iORFs because translated iORFs can be under-detected with existing methods, and we wished to construct a more inclusive list at the expense of some false positives. To do so, we performed a permissive reanalysis of translation initiation site sequencing (TI-seq) data to identify candidate sites of internal translation initiation (see Methods for details and Table S2 for the list). Of 70 iORF candidates identified in the TI-seq analysis, 25 were already represented in Tiers 1 and 2 and the remaining 43 fall into Tier 3.

Importantly, 51 of 70 candidate iORFs can be mapped to alternative transcripts lacking the CDS start codon that are annotated in NCBI and/or GENCODE, including 4/6 Tier 1 iORFs, 17/21 Tier 2 iORFs, and 30/43 Tier 3 iORFs. These alternative transcripts arise from both alternative splicing and alternative transcription start site (TSS) usage (Fig. 1C and table S1), and we highlight several examples to illustrate these mechanisms. As previously reported, exclusion of cassette exon 2 in the *DEDD2* gene skips the CDS start codon to give rise to an iORF-encoding alternative transcript (Fig. 1D). Another example of alternative splicing occurs in *RUSC1*. In this case, an alternative 5′ splice donor site within *RUSC1* exon 1 can be joined directly to exon 3, skipping exon 2 along with the CDS start codon (Fig. 1E). Among the transcripts utilizing alternative TSS, we observed two types. The first type utilizes alternative TSSs within the same first exon but positioned after the start codon of the CDS. An example of this type of iORF can be found in the *GLA* gene (Fig. 1C and Fig. S1). The second type involves alternative TSSs with a novel first exon that bypasses the start codon for the CDS. An illustration of this type of iORF is evident in the *CENPM* gene (Fig. 1C). We conclude that iORF-encoding alternative transcripts that lack the annotated CDS start codon can in many cases be identified in transcriptomic resources.

We next sought evidence for protein-level expression of 20 selected iORFs that can be assigned to annotated alternative transcripts. While expression of Tier 1 and Tier 2 iORFs are supported by translation and HLA I immunopeptidomics, both of these represent indirect evidence and cannot directly infer expression of a stable, cellular microprotein. We utilized translation reporters consisting of alternative transcripts inclusive of the 5′ UTR through the stop codon of the 3′ epitope-tagged iORF. Overexpression followed by immunoblotting or immunofluorescence demonstrated that 14 of the 20 tested iORFs produced microproteins (Fig. S2). Furthermore, CRISPR–Cas9–mediated epitope-tag knock-in at the endogenous *DEDD2* and *RUSC1* loci, combined with immunoblotting, confirmed that these two iORFs are expressed endogenously (Fig. 1F,G). These data suggest that some iORFs arising from annotated alternative transcripts have the potential to generate detectable microproteins.

We next tested the hypothesis that iORFs are preferentially translated from alternative transcripts. We cloned four selected iORFs along with the corresponding upstream sequences from both the annotated canonical transcripts, which contain the upstream CDS start site, and alternative transcripts, which do not, into an overexpression vector, incorporating an epitope tag reporter at the 3′ end of the iORF. Translation of the *DEDD2, CENPM*, and *GLA* iORFs was exclusively observed from the respective alternative transcripts and not the canonical transcripts containing the upstream CDS start sites (Fig. 2A). While the canonical *RUSC1* transcript supported detectable iORF translation, the alternative transcript supported dramatically higher microprotein levels. Consistent with the scanning model of translation initiation, when we mutated the CDS start codons within the canonical transcripts for *RUSC1* and *GLA* (selected because only one AUG start codon is present before the iORF in both of these transcripts), iORF translation became detectable (Fig. S3A). We also confirmed that the annotated CDS can be expressed from each canonical transcript, as expected (Fig. S3B). We conclude that alternative transcripts can be required to bypass translational repression by the start codon of the CDS and enable expression of iORFs.

**Fig 2.**
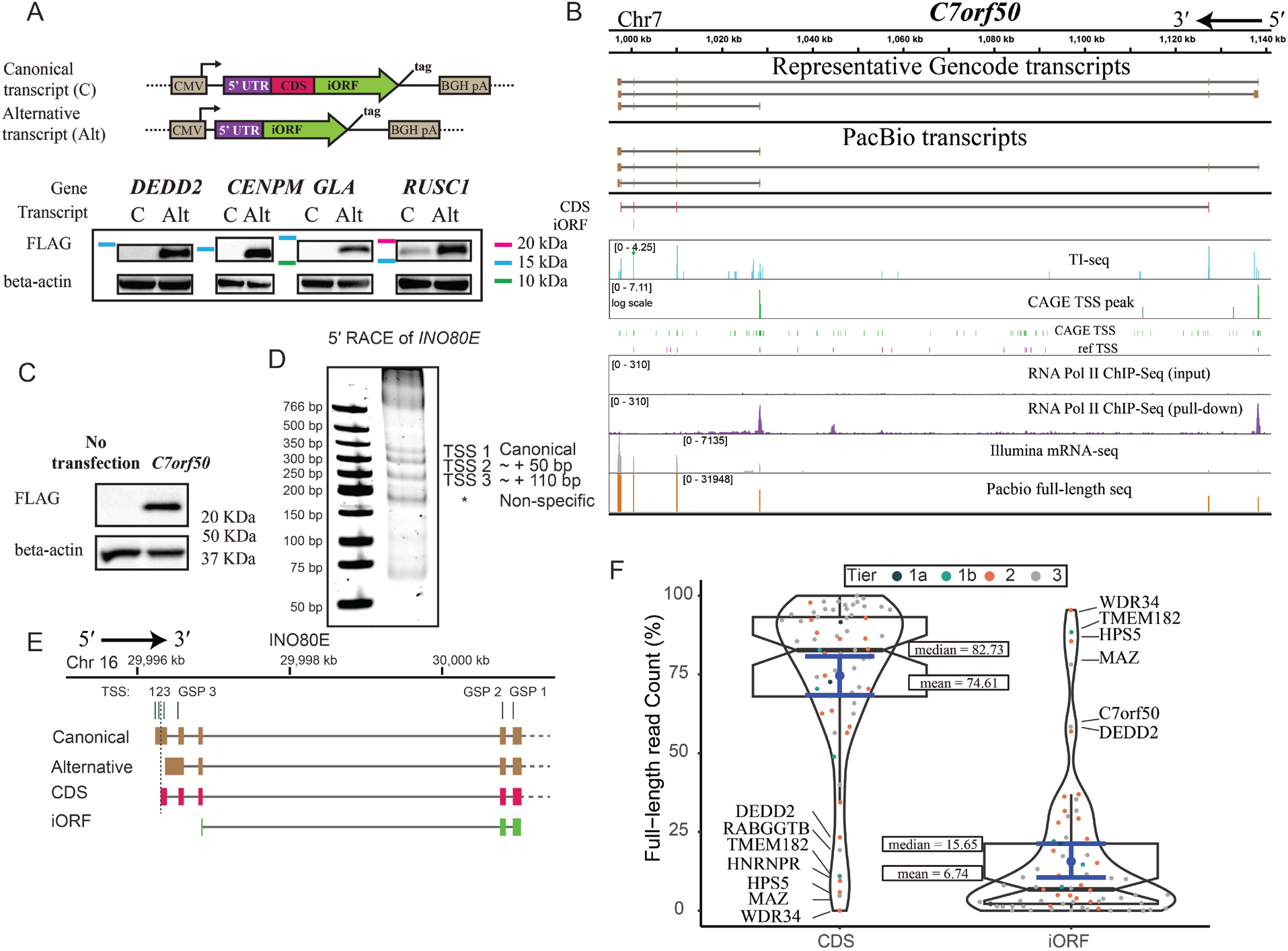
Evidence, classification, and distribution of iORF-encoding transcripts in target genes. (**A**) GENCODE canonical (C) and alternative (Alt) transcripts from four representative genes were cloned from HEK 293 cell cDNA from the 5’ end through the iORF stop codon, with an HA-FLAG epitope tag appended to the 3’ end of the iORF. Anti-FLAG Western blotting was performed after transient transfection to probe iORF expression from each context. (**B**) Evidence for alternative transcripts encoding the *C7orf50* iORF. The C7orf50 CDS (magenta) and iORF (green) are depicted below. TI-seq (cyan), CAGE TSS (green/pink), and Pol II ChIP-seq (purple) reveal internal TSS peaks (red arrows) supporting the iORF-encoding alternative transcript. (**C**) The *C7orf50* alternative transcript (5’ to stop codon of iORF, with a FLAG tag appended to the 3’ end of the iORF sequence) was cloned into an expression vector and transiently transfected into HEK 293T cells, with untransfected cells as a control. Anti-FLAG Western blotting was performed to assess translation of the *C7orf50* iORF. (**D**) Nested PCR amplification of 5′ RACE products for *INO80E*. (**E**) Genome browser view showing the mapped transcription start sites (TSSs) and the positions of the gene-specific primers (GSPs) used in the 5′ RACE of *INO80E*. GSP, gene-specific primer used for 5′ RACE. (**F**) Violin plot comparing the relative abundances of canonical CDS-encoding (left) and alternative iORF-encoding (right) transcripts from 69 target genes detected by ORF capture and PacBio sequencing. Data points are color-coded by confidence tier, with median and mean values indicated. Genes exceeding 50% read counts for an iORF-encoding transcript are labeled.

### Validation and quantitation of iORF-encoding alternative transcripts

For annotated iORF-encoding alternative transcripts to be biologically relevant, these transcripts should exhibit expression comparable to the levels of full-length transcripts encoding annotated protein CDS. Furthermore, we hypothesized that for the 70 candidate iORF-encoding genes in our query set, some could exhibit alternative transcripts that are inconsistently or incompletely annotated across resources; conversely, a subset of iORFs may be expressed via leaky scanning downstream of the annotated CDS start site, and the corresponding genes would be expected to exhibit only transcripts containing the complete annotated CDS. We therefore sought to deeply map and quantify all transcripts encoding the 70 candidate iORFs in human cells. To do so, we enriched cDNA corresponding to the genes of interest followed by PacBio sequencing of full-length transcript isoforms(*19, 20*). After transcript isoform deconvolution and quantitation followed by automated identification of transcripts as iORF or CDS-coding (see Methods), we further classified alternative transcripts arising from alternative splicing or alternative TSS usage (Table S1). Where necessary we utilized Cap Analysis of Gene Expression (CAGE) sequencing(*21*) and RNA polymerase II chromatin immunoprecipitation (ChIP) sequencing data(*22*) to provide orthogonal evidence for alternative TSSs.

This analysis validated existing NCBI or GENCODE annotation of alternative transcripts encoding iORFs, and in some cases identified novel iORF-encoding alternative transcripts arising from the same loci. For instance, PacBio sequencing, along with CAGE-Seq and RNA Pol II ChIP-Seq, confirmed the existence of iORF-encoding alternative transcripts that use a downstream alternative TSS in the gene *C7orf50* (Fig. 2B). However, the most abundant alternative transcripts in our dataset have different exon combinations at the 3′ end compared to the annotated variants; while this has no impact on their iORF coding capacity, alternative 3′ termini could alter post-transcriptional regulation and stability. The *C7orf50* iORF falls into class 3, lacking validated Ribo-seq or proteomic evidence for expression. We therefore cloned the *C7orf50* alternative transcript into an expression vector with an epitope tag fused to the 3′ end of the putative iORF sequence in order to provide evidence for protein-level expression.

Interestingly, we observed expression of a ∼20-kilodalton protein in the iORF reading frame, substantially larger than the predicted ∼3-kilodalton product of AUG-initiated translation, possibly suggestive of use of an upstream, in-frame non-AUG initiation site or other well-documented causes of aberrant SDS-PAGE migration such as charge, intrinsic disorder or post-translational modification (*23-26*) (Fig. 2C).

In another example, we validated the existence of an annotated alternative transcript and also identified a novel iORF-encoding transcript utilizing a downstream TSS in the *RPS8* gene (Fig. S4A). While the alternative 5′ transcript end supporting iORF translation that we detect is annotated in GENCODE, it is absent from NCBI, providing a clear example of how inconsistent or absent transcript variant annotation can lead to mis-mapping of iORFs to canonical transcripts (Fig. S4A). *RPS8* is another Tier 3 gene lacking prior evidence for iORF translation; immunoblotting and immunofluorescence of the alternative transcript cDNA supported translation of a <20-kilodalton iORF-encoded protein which again migrates at an apparent molecular weight much higher than predicted (Fig. S4B-C). Overall, we conclude that alternative transcripts may generally support iORF translation, despite their inconsistent annotation status across resources.

We next considered 18 putative iORF-encoding genes that did not exhibit previously annotated alternative transcripts in either NCBI or GENCODE. These genes span all levels of evidence for iORF translation, with 2 Tier 1 iORFs, 4 Tier 2, and 13 Tier 3. In 17 cases, reads that could correspond to alternative transcripts with novel 5′ ends were detected; however, none of these putative transcript ends could be unambiguously assigned to a distinct TSS on the basis of weak or diffuse CAGE-seq or RNA Pol II ChIP-seq signal - likely the reason that they have not previously been detected. To assess whether these represent *bona fide* novel alternative transcripts arising from polydisperse transcription initiation, and exclude the possibility that they are experimental artifacts resulting from premature reverse transcription termination during PacBio library preparation, we performed 5′ rapid amplification of cDNA ends (RACE) and Sanger sequencing on *INO80E*, a gene harboring a high-confidence Tier 1a iORF and putative novel downstream TSSs detected in our PacBio sequencing (Fig. S5). Our 5′ RACE experiment detected a transcript arising from the annotated *INO80E* TSS, as well as two downstream transcript 5′ ends that precisely correspond to the PacBio-identified 5′ end sequences (Fig. 2D and 2E). While the TSS we detect are close in sequence space (within 110 nucleotides), because the annotated *INO80E* start codon is close to the 5′ end of the canonical transcript, the novel TSS are sufficiently downstream to generate alternative transcripts that exclude the canonical start codon and thus only encode the iORF. Although this analysis cannot completely rule out leaky scanning or dual translation of the *INO80E* iORF from the canonical transcript, it validates the existence of novel TSSs at this locus and underscores the incomplete nature of current gene annotations. Similar experimental validation for putative novel TSS within the remaining 17 genes in this group will be required to demonstrate their biological significance, but the example of *INO80E* demonstrates that previously unannotated alternative transcripts have the potential to support iORF expression from these genes.

Finally, we quantified the iORF-encoding alternative transcripts observed in our experiments (including those lacking orthogonal supporting evidence) and compared their abundances to canonical, CDS-encoding transcripts from the same genes in the HEK 293 cells under study (Fig. 2F). In several instances, the alternative transcript emerged as the predominant gene product; importantly, the iORFs encoded in these abundant alternative transcripts span all 3 tiers, including Tier 1b (*TMEM184*), Tier 2 (*DEDD2, WDR34* and *HPS5*), and Tier 3 (*MAZ* and *C7ORF50*). We note that, while the highest-abundance alternative transcripts span iORF translation-evidence tiers, annotated iORF-encoding alternative transcripts could be identified for all of these genes; putative alternative transcripts lacking prior annotated evidence were universally detected at low levels (at or below 1% of reads). While low-abundance alternative transcripts do not necessarily preclude their presence in cells or necessity for iORF expression, we posit that iORF-encoding alternative transcripts exhibiting comparable expression to CDS-encoding transcripts are most likely to be biologically functional.

### Properties of iORF-encoded microproteins

Because iORFs encode microproteins with complete different amino acid sequences than the canonical proteins that they overlap, yet arise from the same DNA sequence, it is important to query their functions experimentally. We selected four Tier 1a iORFs for which we provided the strongest evidence for translation from alternative transcripts (mapping to the genes *DEDD2, CENPM, GLA* and *RUSC1*) and identified their interactomes when overexpressed in human cells.

None interacted directly with the canonical proteins that they overlap, but each significantly enriched a unique set of proteins or complexes when subjected to co-immunoprecipitation and quantitative proteomics (Fig. 3A). Specifically, the *DEDD2* iORF-encoded microprotein enriched a number of mitochondrial proteins, consistent with its previously reported mitochondrial localization(*14*); the *GLA* iORF microprotein enriched the endosome-associated CLINT1 protein with the greatest statistical significance; the *RUSC1* iORF microprotein enriched many members of the T-complex protein Ring Complex (TRiC) complex; and two protein phosphatase 2 regulatory subunits exhibited the largest magnitude of enrichment by the *CENPM* iORF microprotein. Each of these interactomes is furthermore distinct from the previously reported functions of the corresponding canonical proteins (*27-31*). Consistent with their putative interactomes, each iORF microprotein exhibited a unique subcellular localization when overexpressed, which was distinct from that of the corresponding co-encoded, canonical protein (Fig. 3B). We conclude that, consistent with prior reports characterizing iORFs, the sequences, localizations, and interactions of frameshifted microproteins are generally completely distinct from the overlapping protein. This demonstrates that, at least in these selected examples, human genes can encode two different proteins with different molecular and cellular roles via alternative transcripts.

**Fig 3.**
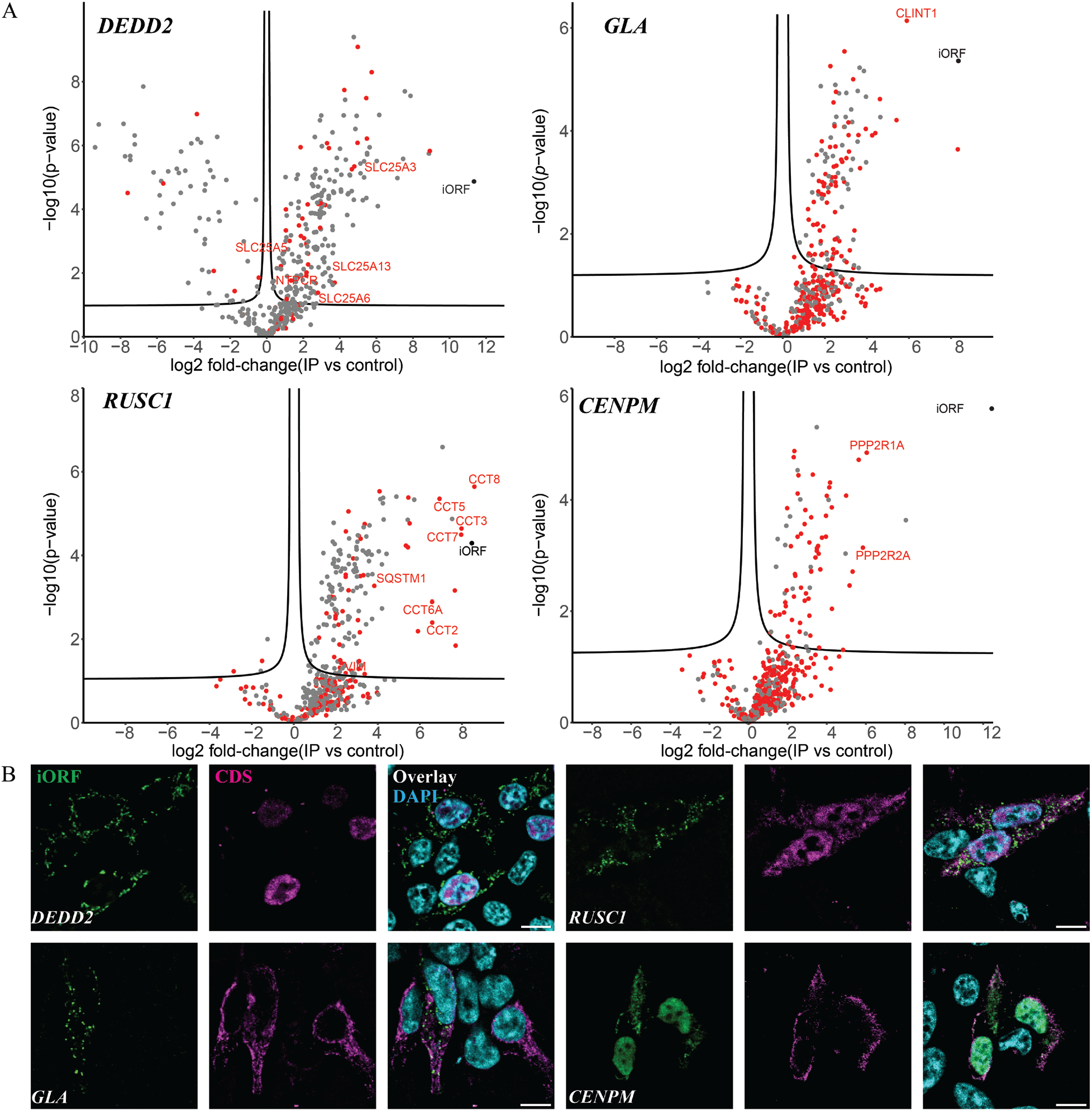
iORF-encoded microproteins have distinct interactions and subcellular localizations. (**A**) Volcano plots from label-free quantitative proteomics of selected FLAG-tagged iORFs co-immunoprecipitated from HEK 293T cells in comparison to controls; threshold, 5% FDR. Enriched proteins annotated (UniProt) as colocalizing to the same subcellular compartment in which the corresponding iORF is predicted to reside are highlighted in red and selected examples are labeled. (**B**) Immunofluorescence of FLAG-tagged iORFs (green) and corresponding HA-tagged canonical proteins (magenta) co-expressed in HEK 293T cells reveals distinct subcellular distributions; DAPI (cyan). Scale bars, 10 µm.

### Regulation and function of the human DEDD2 iORF

In order to examine the cellular consequences of an alternative splicing-driven switch between CDS vs. iORF expression, we selected the Tier 2 *DEDD2* gene. The *DEDD2* iORF was first reported in a peptidomic study (*14*) and has subsequently been supported by high-quality Ribo-seq data (*15*), and we above validated that the iORF-encoding alternative transcript is the major transcriptional output of the *DEDD2* gene in HEK 293 cells (Fig. 2F). Supporting its functionality at the protein level, the *DEDD2* iORF is conserved in higher mammals (Fig. S6A,B). Interestingly, the parental DEDD2 protein is conserved in placental mammals with high amino acid identity except for the iORF-overlapping region, where greater sequence variation is observed (Fig. S6C,D), suggesting that the microprotein-coding iORF may have been overprinted on the DEDD2 CDS.

We next assessed evidence for *DEDD2* alternative splicing in vivo. We reanalyzed RNA sequencing data from 20 cancer cell lines and 20 normal tissues(*32*). The splice junction between exons 1 and 3, which is unique to the alternative transcript encoding the iORF, is detectable in normal human tissues (Fig. 4A, S7A). Reads spanning the iORF-specific exon junction are elevated in cancer cells relative to normal tissue (Fig. 4A). Transcript-level evidence therefore supports DEDD2-SEP expression in normal human tissue as well as in disease.

**Fig 4.**
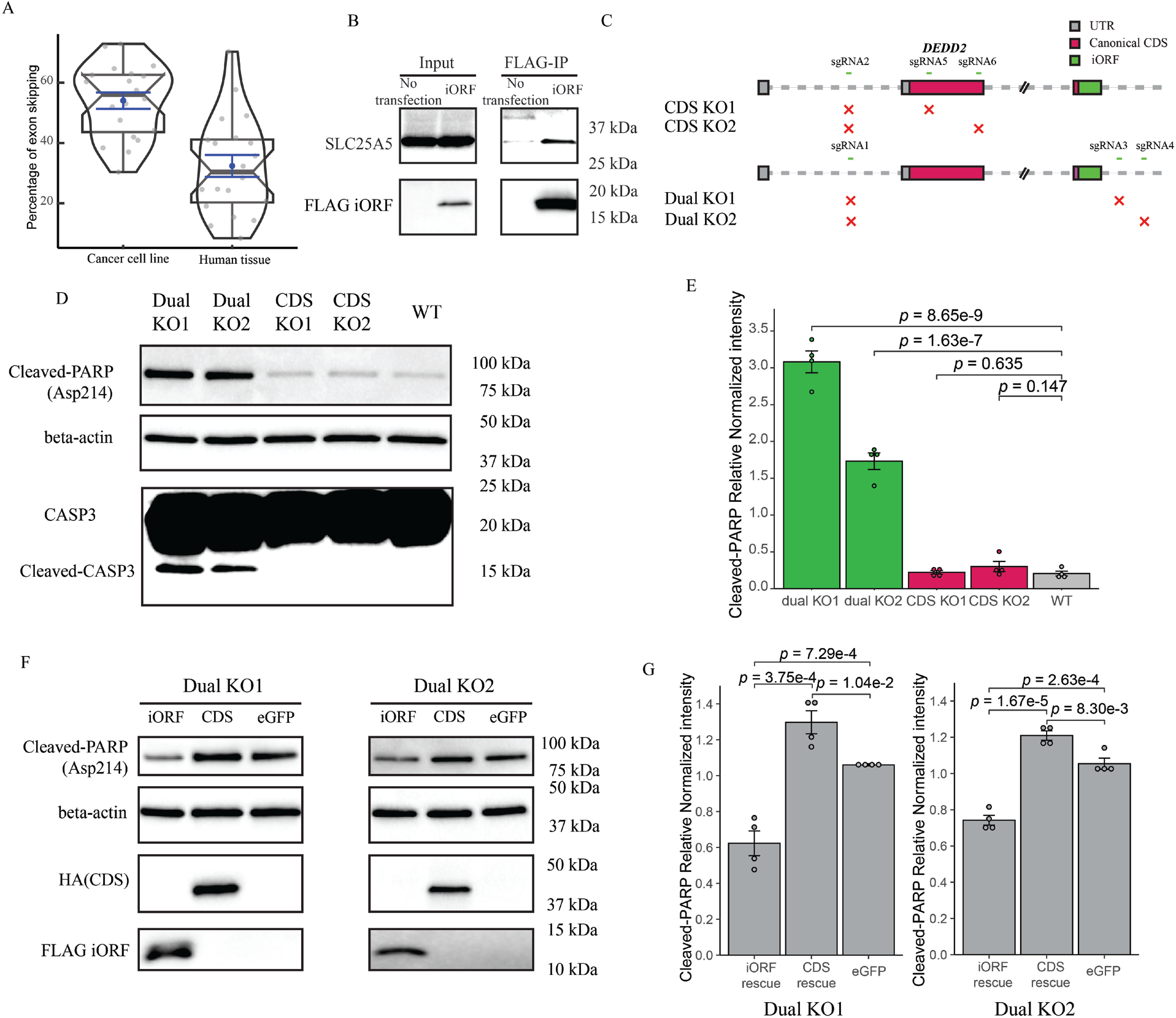
DEDD2 iORF is antiapoptotic. (**A**) Violin plots of ENCODE RNA-seq data showing relative frequencies of DEDD2 exon 2 exclusion (generating the iORF-encoding alternative transcript) in human cell lines and tissues. (**B**) Anti-FLAG co-immunoprecipitates from HEK 293T cells expressing FLAG-tagged DEDD2 iORF, or controls, were subjected to Western blotting with anti-FLAG and anti-SLC25A5 antibodies; lysates were probed as loading controls. (**C**) CRISPR/Cas9 strategy to distinguish the function of the *DEDD2* CDS from the iORF: “CDS KO” deletes exon 2 (CDS-specific), while “dual KO” deletes exons 2 and 3 (shared by CDS and iORF). (**D**) Western blot of cleaved PARP, CASP3, and cleaved CASP3 in WT, dual KO, and CDS KO cell lines with FasL treatment. Data are representative of 4 replicate experiments. (**E**) Quantified PARP cleavage (n=4) from (D). (**F**) Western blotting of cleaved PARP in dual KO cell lysates stably expressing a plasmid encoding either the *DEDD2* CDS, iORF, or eGFP control. (**G**) Quantification (n=4) of PARP cleavage in (F). Bars show mean ± SEM; individual data points represent replicates. All p-values by two-sided Student’s t-test.

Understanding how *DEDD2* alternative splicing is regulated could provide insight into the significance of its variable levels. The RNA-binding protein database(*33*) predicts an SRSF2 binding motif in cassette exon 2 (Fig. S7B). Correspondingly, overexpression of SRSF2 increases the ratio of the canonical transcript that encodes the CDS to the iORF-encoding alternative transcript (Fig. S7C), while knockdown of SRSF2 decreases the relative abundance of the canonical transcript (Fig. S7D). We conclude that SRSF2 binding within exon 2 of the *DEDD2* pre-mRNA promotes its inclusion; because exon 2 contains the annotated CDS start codon, this transcript supports translation of the canonical DEDD2 protein (Fig. S76E). In contrast, decreased SRSF2 binding leads to cassette exon 2 skipping, exclusion of the canonical CDS start codon, and expression of the iORF-encoding alternative transcript.

It is noteworthy that the RNA binding domain of SRSF2 is frequently mutated in myelodysplastic syndromes, chronic myelomonocytic leukemia, and other myeloid malignancies(*34*). While SRSF2 mutations do not always phenocopy absence of SRSF2, we hypothesized that DEDD2 exon skipping and iORF expression could nonetheless be upregulated by cancer-associated SRSF2 mutation. We reanalyzed RNA-seq datasets collected from K562 cells engineered to bear a genomic SRSF2 P95L mutation(*35*), and observed upregulation of the DEDD2 iORF-encoding alternative transcript in these cells relative to wild-type controls (Fig. S7F,G). *DEDD2* pre-mRNA alternative splicing is thus under control of SRSF2 and the DEDD2-SEP-encoding alternative transcript is upregulated by a disease-associated SRSF2 mutation.

Given accumulating evidence supporting functionality of the iORF-encoded DEDD2-SEP microprotein, we sought to characterize its cellular role. Despite their entirely different amino acid sequences, several prior studies have linked the pathways and phenotypes associated with iORFs either positively or negatively to those of the annotated CDS. The annotated DEDD2 protein is homologous to DEDD, a death effector domain-containing protein that is thought to transmit apoptotic signaling into the nucleus (*36*). Like DEDD, DEDD2 has been reported to weakly promote death receptor-dependent apoptosis when it is overexpressed in human cell lines(*27*). We therefore hypothesized that the iORF-encoded DEDD2-SEP could function in apoptosis, and, based on its upregulation in cancer, could oppose the pro-apoptotic function of DEDD2.

To test the role of the DEDD2-SEP microprotein in apoptosis, we first sought to validate an interaction partner (Fig. 3A). We reduced our list of DEDD2-SEP-enriched proteins to those within the mitochondrion (Fig. 3A, red), and noticed that several transporters of the inner mitochondrial membrane (SLC25 protein family) were represented. While it did not exhibit the highest fold-change in our current dataset, across replicate experiments, SLC25A5 was enriched by DEDD2-SEP pull-downs with statistical significance (data not shown), suggesting that this interaction is reproducible. We further utilized AlphaFold3(*37*) to query whether proteins enriched by co-immunoprecipitation are predicted to form plausible complexes with DEDD2-SEP. While we do not seek to validate this predicted model, AlphaFeld3 predicted a low-confidence interaction of an alpha-helical region of DEDD2-SEP with the transmembrane domain of SLC25A5 (Fig. S8A), while several other likely nonspecific enriched proteins do not (data not shown). We therefore validated the interaction of DEDD2-SEP with SLC25A5. Co-immunoprecipitation of FLAG-tagged DEDD2-SEP from human cells followed by Western blotting confirmed robust enrichment of SLC25A5 by DEDD2-SEP, consistent with an association between these proteins (Fig. 4B). SLC25A5 encodes adenine nucleotide translocase-2 (ANT2), an ADP/ATP translocase of the inner mitochondrial membrane. Importantly, the antiapoptotic role of ANT2 is well-characterized (*38*), and, while the molecular mechanism of DEDD2-SEP remains undefined, its interaction with ANT2 is consistent with an antiapoptotic function of the microprotein.

To directly probe the function of the DEDD2-SEP microprotein in human cells, we generated two knockouts (KOs) to differentiate the function of the CDS and iORF: an exon 2 KO abrogating expression of only the canonical CDS, and an exon 3 KO interrupting both the CDS and iORF (Fig. 4C). We then examined cleavage of poly (ADP-ribose) polymerase (PARP) and caspase-3 in these cell lines as a readout of apoptosis downstream of Fas ligand (FasL)(*39, 40*) treatment. While the loss of the CDS alone in two independent clonal cell lines had no significant effect on the cell’s response to FasL, the loss of both the CDS and the iORF, again in two independent clones, greatly increased the level of cleaved PARP and caspase-3 (Fig. 4D,E, and additional clonal cell lines in Fig. S8B), consistent with increased apoptosis in the dual KO. Moreover, dual KO, but not CDS-only KO, significantly slowed cell proliferation (Fig. S8C). These experiments suggest that the cellular phenotype associated with the *DEDD2* gene is suppression of apoptosis – the opposite of the expected result if the *DEDD2* gene is pro-apoptotic as annotated – and that deletion of the exon specific to the annotated DEDD2 protein alone has little, if any, effect on apoptosis.

To fully separate the functions of DEDD2 and DEDD2-SEP, we reintroduced either DEDD2 or DEDD2-SEP into the dual KO cells. Stable reintroduction of the DEDD2 CDS on a plasmid slightly increased the level of cleaved PARP in two independent dual KO cell lines, consistent with previous reports that overexpressing the CDS of DEDD2 weakly promotes apoptosis(*27, 28*) (Fig. 4F,G) but failing to account for the apparent antiapoptotic phenotype observed in the *DEDD2* dual KO. In stark contrast, stable rescue with a plasmid encoding the DEDD2 iORF dramatically decreased FasL-dependent apoptosis in the two dual KO cell lines (Fig. 4F,G).

Taken together, these experiments demonstrate that the iORF-encoded DEDD2-SEP microprotein is antiapoptotic and furthermore predominates the phenotype associated with the *DEDD2* gene in cultured human cells.

## Conclusion

With the prevailing paradigm that same-strand, frameshifted, overlapping ORFs are unique to viruses, coupled with the difficulty in explaining human iORF translation downstream of canonical, CDS initiation sites, human iORFs have generally been overlooked for functional study. We demonstrate in this work that, for iORFs that cannot be translated from canonical transcripts that encode an overlapping CDS, alternative transcripts lacking the CDS start site are required for iORF translation. Due to the incomplete or absent annotation of some of these alternative transcripts, iORFs have been mis-mapped within canonical transcripts encoding the CDS, but in many cases are not translated from this context. Other classes of dual coding genes, including mRNAs containing upstream ORFs, have been shown in some cases to exhibit alternative transcripts (*41*), suggesting that this phenomenon may not be specific to iORFs.

Transcript-vs. translational-level control of iORF expression likely has not only mechanistic, but also functional and regulatory implications. In the case of iORFs that can be co-expressed from the same transcript as the CDS, several studies have suggested that the uORF-encoded protein and downstream protein have related functions or phenotypes, and in some cases may even directly interact(*3, 8, 42*). In contrast, iORFs that are expressed from alternative transcripts can be expressed under distinct conditions as the CDS, and thus can have different or even opposing functions (as in the case of *DEDD2*). To determine whether this model generalizes to more iORF-encoding genes will require further identification of iORF vs. CDS-encoding transcripts, as well as experimental investigation of iORF functions. It will also be critical to determine if dual translation vs. alternative transcript production differentiates functional outcomes for other classes of dual-coding genes, including upstream and downstream ORFs, as well.

An important unanswered question raised by the existence of iORFs is the mechanism by which their overlapping sequences arise in evolution. While we do not attempt to address this question in our study, we can consider whether and how our results comport with models of *de novo* gene origination and overprinting, since understanding iORF molecular evolution will be just as critical as elucidating their expression mechanisms to support their biological reality and functionality. While acquisition of frameshifted iORFs within viral genes is known, it is nonetheless somewhat surprising that this process might occur in human genes, as mutations required to create a start codon in the iORF reading frame could be deleterious in the CDS reading frame. *De novo* gene origination from non-coding regions (*43, 44*) is not subject to this limitation. We observed in this work that the overlapping region shared by the *DEDD2* CDS and iORF exhibits lower sequence conservation in the CDS reading frame than the rest of the protein, contrasting the naïve expectation that the shared region might be more evolutionarily constrained than the rest of the CDS by the presence of functional coding sequences in two reading frames.

We speculate that the DEDD2 iORF could have recently arisen within a region of the *DEDD2* CDS that tolerates mutation, generating two functional, overlapping proteins; alternatively, it is possible that the evolutionarily duplicated, non-essential DEDD2 protein is undergoing pseudogenization; a mutation creating the *DEDD2* iORF would then be more analogous to *de novo* gene origination within a non-coding RNA. These observations interestingly parallel our recent report that the *E. coli* GndA-encoding iORF overlaps a less-conserved region of the coding sequence for 6-phosphogluconate dehydrogenase (*45*). Expanded and rigorous functional and evolutionary analysis will be required to correlate iORF evolution, expression mechanisms and function more broadly.

Regardless of their origins and expression mechanisms, the existence of iORFs that overlap canonical protein CDSs has substantial implications for understanding and treating disease(*42*). Disease-associated mutations mapping to genomic regions encoding overlaps, especially for the genes identified in this study,(*46, 47*) should be reconsidered to determine if the iORF contributes to the disease phenotype, as has previously been suggested for the alt-FUS and alt-RPL36 overlapping proteins. The presence of iORFs should also be carefully considered when designing siRNA therapies, because iORF-encoding alternative transcripts share most of their sequence with canonical transcript targets.

## Acknowledgments

This work was supported in part by the National Institutes of Health (1R01GM155404), an Emerging Leader Award from the Mark Foundation for Cancer Research, a Distinguished Investigator Award from the Paul G. Allen Frontiers Group, and a Sloan Research Fellowship (FG-2022-18417), to S.A.S. We thank members of the Slavoff and Loh labs for helpful discussions. We also thank the Yale West Campus Analytical Core (WCAC) for their assistance with LCMS. We thank Yale Center for Genome Analysis and Keck Microarray Shared Resource at Yale University for providing PacBio sequencing and high-performance computing services, which are funded in part by the National Institutes of Health instrument grants 1S10OD028669-01 and 1S10OD030363-01A1, respectively. We thank the Yale West Campus Imaging Core for providing confocal microscopy.

## Author contributions

Conceptualization: H.S., S.A.S., S.G.K.

Methodology: H.S., S.A.S.

Software: H.S.

Investigation: H.S., S.A.S.

Visualization: H.S., S.A.S.

Funding acquisition: S.A.S.

Project administration: H.S., S.A.S.

Supervision: S.A.S.

Writing – original draft: H.S., S.A.S.

Writing – review & editing: H.S., S.A.S., S.G.K.

## Data and materials availability

The PacBio sequencing data are deposited in the Sequence Read Archive with accession number PRJNA1130103. MS-based proteomics data are available via PRIDE with accession number PXD053420. During review, log in to the PRIDE website using the following details:

Username

Password: xOKh0QreU491

## Supplementary Materials

Materials and Methods Figs. S1 to S8

Tables S1 to S4

