## Supplementary Information for "Nested small open reading frames are translated from alternative transcripts"

Science
AAAS

Supplementary Materials for

### Nested human small open reading frames are translated from alternative transcripts

Haomiao Su,[1,2] Samuel G. Katz[3], Sarah A. Slavoff[1,2,4,*]

**The PDF file includes:**

Materials and Methods
Supplementary Text
Figs. S1 to S8

#### Materials and Methods

##### Antibodies

The following primary antibodies were used for immunoblotting: anti-FLAG (1:1000, Sigma, F1804), anti-HA (1:3,000, Invitrogen, 71-5500), anti-β-actin (1:3,000, Invitrogen, MA5-15739), anti-SLC25A5 (1:1,000, Invitrogen, MA5-15739), anti-Histone-H3 (1:2000, Cell Signaling, 4499), anti-Cleaved-PARP (Asp214) (1:1000, Cell Signaling, 5625). The following secondary antibodies were used for immunoblotting: goat anti-rabbit IgG horseradish peroxidase conjugate (1:4,000, Rockland, 611-1302) and goat anti-mouse IgG horseradish peroxidase conjugate (1:4,000, Rockland, 610-1319). Primary antibodies for immunostaining were mouse anti-FLAG (1:1000, Sigma, F1804) and rabbit anti-HA (1:500, Invitrogen, 71-5500). Secondary antibodies for immunostaining were goat anti-rabbit IgG Alexa Fluor 568 (1:500, Invitrogen, A11011), goat anti-rabbit IgG Alexa Fluor Plus 555 (1:500, Invitrogen, A32732), goat anti-mouse IgG Alexa Fluor 647 (1:500, Invitrogen, A21235), and goat anti-mouse IgG Alexa Fluor Plus 647 (1:500, Invitrogen, A32728).

##### Assignment of iORF Tier

To assign confidence tiers to iORFs, we compiled ribosome profiling (RIBO-seq) evidence from GENCODE's gold-standard human translatome resource (*15*) and peptide-spectral match (PSM) data from a community reanalysis of proteomics datasets deposited in PeptideAtlas(*18, 48*). Only PSMs that passed manual quality checks were considered valid evidence. Specifically, we included only non-HLA PSMs ranked 1 or 2, and HLA-associated PSMs with positive manual annotations.

Tier assignments were made based on the strength of combined RIBO-seq and mass spectrometry (MS) support. **Tier 1a** iORFs are the highest-confidence candidates, supported by curated RIBO-seq data and at least two unique PSMs from PeptideAtlas. **Tier 1b** iORFs are supported by RIBO-seq and at least one unique, curated PSM. **Tier 2** iORFs are supported solely by curated RIBO-seq evidence. These criteria ensure that higher-tier iORFs are backed by multiple, independent lines of translational evidence. **Tier 3** ORFs are those not reported in the GENCODE study but identified by TI-seq.

##### Identification of anisoforms

To identify candidate anisoforms, lactimidomycin (LTM)-based translation initiation site sequencing (TI-seq) data (SRA: SRR618772 and SRR618773) generated from HEK293 cells were utilized. Adaptor trimming was performed using Cutadapt (version 4.4), and transfer RNA and ribosomal RNA were filtered out with STAR (version 2.7.11a). The remaining reads were aligned to the hg38 reference genome with the guidance of GENCODE Human Release v38 using STAR. Subsequently, the mapped reads were analyzed using PRICE (Gedi version 1.0.2) with the assistance of Ensembl annotation version 104 to identify codons. The resulting cit file was then converted to bedgraph format using the ViewCIT command. A python script (TI_Seq_ORF.py, available at https://github.com/slavofflab/TI-seq-ORF) was used to process the bedgraph files. A peak, defined at the codon level, was called when satisfied following conditions: (i) The peak is among the top 5 highest signals in the gene. (ii) The peak intensity reached 5 observed reads calculated by PRICE. (iii) The peak intensity is greater than 20% of the highest intensity in the gene. (iv) The position must be a local maximum within a span of seven nucleotides. We subsequently assigned peaks to ORF types based on their positions within GENCODE V38 transcripts. The types include ncORF (ORF within non-coding RNA), CDS (annotated coding sequence), N-extension (unannotated N-terminal extension in-frame with annotated protein CDS), uORF (ORF upstream of an annotated CDS), uoORF (ORF that initiates upstream and partially overlaps an annotated CDS in an alternative reading frame), ooORF(outside overlapping ORF that initiates before, ends after, and entirely encompasses the annotated CDS in an alternative reading frame), N-deletion (N-terminal truncation/internal, in-frame translation initiation site of an annotated CDS), dORF (ORF downstream of an annotated protein CDS), doORF (ORF that initiates within CDS, and partially overlaps an annotated CDS in an alternative reading frame), and iORF (internal, frameshifted, overlapping ORF within an annotated CDS). We then implemented a computational procedure to identify candidate iORFs that may be expressed from either known or unannotated anisoform alternative transcripts – in other words, anisoforms. First, we rigorously identified a training set of iORFs with evidence for alternative transcripts in GENCODE V38. If a peak is designated as an iORF or doORF in any transcript isoform and is not assigned to CDS or any of their isoforms in any transcript isoform the peak will be considered as a candidate for follow up study. Only iORFs with AUG start codons were selected for validation. The first round of anisoform candidates were then manually checked against the following criteria with the order from highest to lowest intensity: (1) expression of the annotated alternative transcript for an anisoform that is caused by exon skipping, alternative 5' splicing site and/or alternative first exon should be supported by mRNA-Seq data. Specifically, exon skipping must be evidenced by reads spanning the splice junctions. Similarly, the alternative 5' splice site and the alternative first exon should have reads mapped to their respective specific regions. The mRNA-Seq data (SRA: SRR24971804, SRR24971805, SRR24971806, SRR24971813, SRR24971814, SRR24971815) were aligned to the hg38 reference genome using the GENCODE Human Release v38 annotation as a guide, employing STAR after trimming and filtering out non-coding RNA reads. The resulting bam files were merged by samtools (version 1.18) to get the final single bam file. (2) The anisoform should exhibit other in-frame Ribo-Seq peaks. The Ribo-Seq (SRR618771) data underwent identical preprocessing steps for TI-Seq analysis. The Ribo-Seq data (SRR618771) underwent identical preprocessing steps as those used for TI-Seq analysis. No ORF calling was applied. Instead, the observed reads, calculated by PRICE, were used to manually check for in-frame peaks. The in-frame peaks of the iORF should appear at a comparable level (>10% of adjacent in-frame peaks of the CDS), and the in-frame peaks of the third ORF should be absent or present at a much lower frequency. This helps to ensure that the in-frame peaks of the iORF are not produced by non-specific signals. (3) If the annotated alternative transcript for an anisoform uses an alternative transcription start site (TSS), it should be supported by the refTSS database(*49*) or CAGE-Seq or RNA Pol II-Seq. The CAGE-Seq results were downloaded directly from FANTOM5 with FF ontology id 10450-106F9. The RNA Pol II ChIP-Seq data (GSE152062) were aligned to human hg38 genome with STAR after trimming. Next, we identified a second set of experimental iORFs for further validation at the mRNA level. All selection criteria were followed, except for the requirement of existing annotation.

#### Cloning

The constructs pcDNA3.1-HA, pcDNA3.1-FLAG, and pcDNA3.1-HA-FLAG were created by introducing HA, FLAG, and HA-FLAG tags into pcDNA3.1, respectively, using *Eco*RI and *Xba*I restriction enzymes. To validate each anisoform, total RNA was extracted from HEK293T cells using TRIzol (Invitrogen, 15596026), followed by cDNA synthesis with the cDNA Synthesis Kit (NEB, E6300S). Target alternative transcript isoforms (iORF) and canonical transcripts isoforms (CDS) were PCR amplified from the 5' UTR to the stop codon of ORFs. Restriction sites were introduced using gene-specific primers and Q5 polymerase (NEB, M0491S) with High GC Enhancer. The PCR program ran for 35 cycles, each cycle comprising denaturation at 98°C for 10s, annealing at 62°C for 15s, and extension at 72°C for 30s (with 2 mins final extension). The PCR products were purified via agarose gel electrophoresis, digested with corresponding restriction enzymes, and then ligated into pcDNA3.1 vectors containing different tags for various purposes. Following transformation into DH5-alpha cells, single colonies were selected and confirmed by Sanger sequencing to ensure the correct transcript isoform (from 5' UTR to stop codon of selected ORF) was obtained. The *C7orf50* 5' UTR-iORF-HA-FLAG construct was synthesized by GenScript and inserted into pcDNA3.1.

For shRNA lentivirus, the pLKO.1-TRC cloning vector (Plasmid #10878) was digested using *Eco*RI and *Age*I restriction enzymes, followed by purification via agarose gel electrophoresis. shRNA oligos (obtained from Sigma) were annealed in T4 ligase buffer and then inserted into pLKO.1 using T4 ligase. For stable expression, cDNAs were inserted into pLJM1-eGFP (a gift from David Sabatini, addgene plasmid #19319) vectors modified with FLAG/HA tags utilizing *Eco*RI and *Age*I restriction sites. The lentivirus plasmids were transformed into NEB® Stable Competent *E. coli.* Single colonies were selected and confirmed by Sanger sequencing to ensure the correct construction was obtained.

**Cell culture**

HEK 293 (CRL-1573) and HEK 293T cells (CRL-3216) were purchased from ATCC, and early-passage stocks were established to ensure cell line identity; cells were maintained up to no more than 10 passages. Cells were cultured in DMEM (Corning, 10-013-CV) with 10% FBS (Sigma, F0392) and 1% penicillin-streptomycin (Gibco, 15140122) in a 5% CO2 atmosphere at 37 °C.

**Immunofluorescence**

One day before transfection, HEK293T cells were seeded in a 24-well plate with antibiotic-free DMEM with 10% FBS. The cells were 70-90% confluent at the time of transfection. Plasmids were transfected using Lipofectamine 2000 (Invitrogen, 11668019) or Lipofectamine 3000 (Invitrogen, L3000015) according to the manufacturer's instructions. After 6-8 hours, the transfected cells were reseeded onto poly-L-lysine (Sigma, P4832) coated glass coverslips in a 12-well plate in complete media. After allowing the cells to attach overnight, the cells were fixed with 10% formalin (Fisher, SF100-4), quenched with 10 mM glycine in PBST (phosphate-buffered saline with 0.1 % Tween-), and permeabilized with 0.25% Triton X-100 in PBST. After rinsing 3 times with PBST for 5 minutes each, the cells were blocked with 1% BSA in PBST at room temperature for 1 hour. Then, the cells were incubated with primary antibodies (1:300 dilution) in PBST with 1% BSA at 4°C overnight. After rinsing 3 times with PBST for 5 minutes each, the cells were incubated with fluorochrome-conjugated secondary antibodies (1:300 dilution) and DAPI in PBST with 1% BSA for 1 hour, protected from light. After rinsing 3 times with PBST for

5 minutes each, the glass coverslips were mounted onto microscope slides. Confocal imaging was performed on a Leica SP8 LS confocal microscope with a 63× oil immersion objective. The images were processed with ImageJ (version 1.54g) using the DeconvolutionLab2 plugin.

**Immunoprecipitation**

For IP-WB of transfected cells and stable expression cells, the HEK293T cells were grown in T25 flasks. For IP-WB of FLAG KI cells and IP-MS, the HEK293T cells were grown in 15 cm dishes. Cells were transfected using Lipofectamine 2000, Lipofectamine 3000, or polyethyleneimine (Polysciences, 23966) according to the manufacturer's instructions. The cells were lysed with RIPA buffer in the presence of a protease inhibitor cocktail (Roche, 11836170001). After centrifuging at 14,800 rpm for 30 minutes, the supernatant of the lysate was incubated with 15 µl (for T25 flasks) or 30 µl (for 15 cm dishes) of ANTI-FLAG® M2 affinity gel (Sigma, A2220) at 4 °C overnight. After removing the supernatant, the beads were washed with RIPA buffer 3 times. Elution was in 30 µl of 3× FLAG peptide (Sigma, F4799) at a final concentration of 100 µg $ml^{-1}$ in RIPA buffer at 4 °C for 2 h. Eluted proteins were subjected to SDS–PAGE or Tricine–SDS-PAGE for WB or LC-MS/MS.

For co-IP-WB, HEK293T cells were grown in 15 cm dishes and transfected using polyethyleneimine (Polysciences, 23966) according to the manufacturer's instructions. The subsequent steps, including cell lysis, centrifugation, incubation with ANTI-FLAG® M2 affinity gel, washing, elution, and analysis, were performed as described above.

**Proteomics**

Gel slices (protein bands or entire lanes) were digested with trypsin (Promega, V5111) at 37 °C overnight. After reduction by DTT (Sigma, D0632) and alkylation by IAA (TCI, 144-48-9), the resulting peptide mixtures were extracted from the gel and dried. Residual detergent was removed with ethyl acetate followed by desalting with a peptide cleanup C18 spin column (Pierce, 89870). Peptides were resuspended in 35 µl of 0.1% formic acid (FA), followed by centrifugation at 14,800 rpm at 4 °C for 30 min. 5 µl of each sample was injected on a C18 nano column (CoAnn, HEB05005001718I) attached to an EASY-nLC (Thermo) in-line with a Thermo Scientific Q Exactive Plus Hybrid Quadrupole Orbitrap mass spectrometer. A 95-minute gradient was used to further separate the peptide mixtures as follows (solvent A: 0.1% FA; solvent B: 80% acetonitrile (ACN) with 0.1% FA) at a flow rate of 0.1 µl/min: The initial condition was 5% solvent B. Solvent B was then increased to 45% over 60 minutes. The gradient increased to 85% B over the next 1 minute. Solvent B was held at 85% for 10 minutes. Finally, the gradient returned to 5% B over 23 minutes until the end of the run. The full MS spectrum was collected over the mass range of 300–1,700 m/z with a resolution of 70,000, and the automatic gain control target was set at $3 \times 10^6$. MS/MS data were collected using a top 10 high-collision energy dissociation method in data-dependent mode with a normalized collision energy of 27.0 eV and a 1.6-m/z isolation window. MS/MS resolution was 17,500, and dynamic exclusion was 90 s.

The proteomic data were analyzed using MaxQuant (version 2.0.3.0). Oxidation of methionine and protein N-terminal acetylation were set as variable modifications. Carbamidomethylation of cystine was set as a fixed modification. The data were searched against the human UniProt protein database (version 2021) plus the corresponding anisoform sequence. For all analyses, a mass deviation of 20 ppm was set for MS1 peaks, MS/MS tolerance was 0.6 Da, and a maximum of two missed cleavages were permitted. Maximum false discovery rates were set to 1% both on peptide and protein levels. The minimum required peptide length was five amino acids. Protein quantitation was calculated by the label-free quantification and setting the LFQ min ratio count to 2. Data normalization, filtering, and imputation were performed using Perseus (version 2.0.3.0), and the enrichment analysis was performed using the volcano plot with FDR set to 0.05 and S0 set to 0.1. Missing values were imputed from a normal distribution with a downshift of 1.8 and a width of 0.15.

**Construction of CRISPR-Cas9 mediated knock-in and knock-out cell lines**

Guide RNAs (gRNAs) were designed with CRISPOR(*50*) and are listed in Supplementary Table 5. The HDR templates for KI were single-stranded DNA oligonucleotides (ssODN)(*51*) and are listed in Table S3. Double-stranded DNA oligonucleotides corresponding to the gRNAs were inserted into pSpCas9(BB)-2A-GFP vector (a gift from Feng Zhang, Addgene plasmid #48138).

For generation of KO cells, 625 ng of each two gRNA plasmids were co-transfected with 2.5 µg P3000 and 3.75 µg Lipofectamine 3000 (Invitrogen, L3000015) according to the manufacturer's instructions into HEK293 cells in 12-well plate. The top 5% GFP-positive cells were sorted using a BD Aria flow cytometer and seeded into a 96-well plate at a density of <1 cell per well. The genomic DNA from clonal strains was isolated using 0.05% SDS and 16 units/mL of proteinase K (NEB, P8107S) in 10 mM Tris-HCl, pH 8 at 37°C for 2 hours. Subsequently, deactivation was carried out at 80°C for 30 minutes. Target loci were amplified using Q5 polymerase (NEB, M0491S) with GC enhancer and analyzed by agarose gel electrophoresis. PCR products from correctly cloned single cells were purified using spin columns, followed by re-amplification using OneTaq polymerase (NEB, M0480S) with GC Reaction Buffer. Subsequently, they were subcloned using the TOPO™ TA Cloning™ Kit (Invitrogen, 450641) and validated by Sanger sequencing. The generation of knock-in (KI) cells followed the same protocol, except for transfection: 1 µg of gRNA plasmid and 2 µL of 10 µM ssODN were transfected into the cells.

**Generating stable cell lines**

HEK293T cells were seeded in a 6-well plate and transfected with 500 ng of either the shRNA vector or stable expression vector, along with 375 ng of psPAX2 (a gift from Didier Trono, addgene plasmid #12260) and 125 ng of pMD2.G (a gift from Didier Trono, Addgene plasmid #12259) using Lipofectamine 3000. The resulting lentivirus was harvested at 48 and 72 hours post transfection. The combining viral solution was filtered using a 0.45 µm syringe filter (Millipore, SLHA033SS) and used to infect cells in the presence of 8 µg/mL polybrene (Sigma, H8268). Two days post-infection, the cells were subjected to selection with 4 µg/mL Puromycin (Sigma, P8833). Subsequently, the cells were maintained in culture with 2 µg/mL Puromycin.

**Generating ORFeome and biotin-labeling probes**

A fragment between 600 bp to 1.5 kb was amplified from the target genes from cDNA using specific primers and Taq polymerase (NEB, M0480S) with GC Reaction Buffer. The program was run for 45 cycles, with each cycle consisting of denaturation at 94°C for 30s, annealing at 52°C for 15s, and extension at 68°C for 2 min. The PCR products were subcloned directly with TOPO™

TA Cloning™ Kit (Invitrogen™, 450641). Then, biotinylated cDNAs for each clone were synthesized by PCR with universal primers for pCR®2.1-TOPO® vector using a modified dNTP mixture containing 33% biotin-dUTP (prepared by mixing 5 µL of dATP, dCTP, dGTP, 3.35 µL of dTTP (NEB, N0447S) and 33 µL of biotin-16-Aminoallyl-2′-dUTP (Trilink, N-5001)). After purifying with a mini spin column (Econospin, 1910-250), the biotin-labeled cDNAs were randomly fragmented in microTUBEs (Covaris, 520052) on a Covaris S220 sonicator. The sonication method parameters are as follows: peak power of 175 W, duty cycle of 10%, 200 cycles per burst, and duration of 430 s. The fragmented products were dried in a *SpeedVac* (Thermo Scientific Savant SPD1010) and resuspended in $ddH_2O$ to get a high-concentration solution. Finally, the fragmented biotin-labeled cDNAs were mixed to generate the biotin-labeling probe sets at 1 ng/ µL.

**ORF-capture sequencing**

The total RNA isolated from HEK293 cells with TRIzol (Invitrogen, 15596026) was sent to Yale Center for Genome Analysis (YCGA) for library construction utilizing the "Preparing Iso-Seq libraries using SMRTbell prep kit 3.0" (PacBio, PN 102-396-000 REV02 APR 2022) protocol. During the process, the full-length cDNA was taken back after Step 2.4.16. The cDNA was then amplified using NEBNext® High-Fidelity 2X PCR Master Mix (NEB, M0541S) and cDNA primers bc1001-F/R in a 100 µL reaction. The PCR program was run for 14 cycles after initial heating at 98°C for 45 seconds. Each cycle consisted of denaturation at 98°C for 10 seconds, annealing at 62°C for 15 seconds, and extension at 72°C for 3 minutes, with a final extension step at 72°C for 5 minutes. The PCR products were cleaned using 1.8X AMPure XP beads (BECKMAN COULTER, A63880) according to manufacturer's protocol and eluted with 50 µL nuclease-free water.

The amplified cDNA was subjected to target transcript enrichment using the "cDNA Capture Using IDT xGen® Lockdown® Probes" protocol (PacBio, PN 101-604-300 Version 01 June 2018) with modifications. Gen Hybridization™ and Wash Kit (IDT, 413096243) were utilized for this purpose. 1 µg of purified cDNA was combined with 1 nmol of PCR primer bc1001-F/R (Sigma) and 1 nmol of PolyT blocker Oligo (IDT) in a 1.5 mL LoBind tube (Eppendorf, 022431021). After drying in a *SpeedVac* (Thermo Scientific Savant SPD1010), 8.5 µL of 2X Hybridization Buffer, 2.7 µL of Hybridization Buffer Enhancer and 1.8 µL of nuclease-free water were added to resuspend the DNA. The resulting solution was transferred to a PCR tube and cDNA was denatured by placing it in a 95°C Thermal Cycler (Bio-Rad, C1000) for 10 minutes. The tube was quickly spun down after heating. Once the solution cooled to room temperature, 4 µL of biotin-labeling probe sets (4 ng) were added, mixed thoroughly, and then spun down. The mixture was incubated at 65°C for 4 hours. Subsequently,100 µL of M-270 streptavidin beads (Invitrogen) was added and a series of washes were performed according to the protocol. After adding 50 µL nuclease-free water, on-bead PCR was performed in a total volume of 300 µL with the same program used previously, except the cycle number was increased to 30. The PCR product was cleaned using 1.8X AMPure XP according to manufacturer's protocol and eluted with 50 µL nuclease-free water. The cDNA was sent back to YCGA, and the process continued at step 4.1. The final library was sequenced on a PacBio Sequel II.

**Long-read data processing**

The analysis followed the PacBio Iso-Seq3 (https://github.com/ylipacbio/IsoSeq3) workflow. The HiFi reads were initially processed using lima (version 2.7.1) to demultiplex, remove barcode sequences, adjust orientation, and eliminate unwanted primer combinations. Subsequently, the resulting BAM file underwent further refinement with isoseq3 refine (version 4.0.0) to ensure the presence of polyA and to remove concatemers. Following this, the refined full-length reads were clustered using isoseq3 cluster (version 4.0.0). After filtering out low-quality reads, the clustered reads were aligned to the hg38 reference genome using pbmm2 (version 1.13.0). Then, isoseq3 collapse (version 4.0.0) was employed to annotate the transcriptome. The annotated transcripts were sorted using pigeon sort (version 1.1.0), and pigeon classify (version 1.1.0) was utilized to classify and quantify transcripts, guided by GENCODE V38 annotation. Finally, the PacBio transcript isoforms of target genes were classified based on their potential coding abilities with a python script (coding_assign.py, available at https://github.com/slavofflab/Codingtype_assign) based on the first start codon with the mRNA sequence. The RNA-seq datasets mentioned above were aligned to the hg38 genome using Salmon (version 1.10.2). The quantified data was then subjected to further analysis with DESeq2 (version 1.42.1) to obtain the Transcripts Per Million (TPM) values for each gene. The TPM values of transcript isoforms were calculated by multiplying the TPM of the gene with the transcript percentage derived from the long-read sequencing data.

**RT-PCR**

Total RNA was extracted from cells using TRIzol (Invitrogen, 15596026), followed by cDNA synthesis using the iScript™ cDNA Synthesis Kit (BIO-RAD, 1708891). Subsequently, RT-PCR was performed with Luna® Universal qPCR Master Mix (NEB, M3003L) in CFX96 Touch (Bio-Rad, 1854095). The RT-PCR program was initiated with an initial heating step at 95°C for 60 seconds, followed by 45 cycles. Each cycle comprised denaturation at 95°C for 15 seconds and extension at 60°C for 1 minute, along with plate reading. Finally, a melt curve step was conducted from 60-95°C over 30 minutes.

**RAPID AMPLIFICATIONOFcDNAENDS(5'-RACE)**

Total RNA from HEK293T cells was subjected to 5′ RACE using the 5′ RACE System for Rapid Amplification of cDNA Ends (Invitrogen, Cat. No. 18374058) according to the manufacturer's instructions. Gene-specific primers are listed in Table S3. Nested PCR products were then subcloned with the TOPO™ TA Cloning™ Kit (Invitrogen, Cat. No. 450641) and validated by Sanger sequencing.

**Conservation and evolution analysis**

Sequences of DEDD2 and DEDD2-anisoform were obtained from NM_001270614.2 (human, *Homo sapiens*), XM_024351836.3 (Chimpanzee, *Pan troglodytes*), XM_015124083.2 (rhesus monkey, *Macaca mulatta*), XM_008567239.1 (colugo, *Galeopterus variegatus*,), NM_207677.3 (house mouse, *Mus musculus*), XM_003748789.4 (brown rat, *Rattus norvegicus*), NM_001076017.1 (cattle, *Bos taurus*), XM_004015286.5 (sheep, *Ovis aries*), XM_023651436.1 (horse, *Equus caballus*), XM_014867556.2 (donkey, *Equus asinus*), XM_019819482.3 (cat, *Felis catus*), XM_002923941.4 (giant panda, *Ailuropoda melanoleuca*), XM_023544014.1 (elephant, *Loxodonta africana*), XM_023742697.1 (manatee, *Trichechus manatus*), XM_007942940.1

(aardvark, *Orycteropus afer*). Multiple sequence alignment was performed with Clustal Omega (https://www.ebi.ac.uk/jdispatcher/msa/clustalo).

**Statistical analysis**

Statistical analyses were performed using two-sided, two-sample Student's t-test with Microsoft Excel (version 2404) or Welch two Sample t-test with R (version 4.3.3). Error bars represent the mean ± SEM. Further statistical details of experiments are reported in the figure legends. No statistical methods were used to predetermine sample size. The experiments were not randomized, and investigators were not blinded to allocation during experiments and outcome assessment.

**Splicing variation analysis**

For cancer cell lines and tissue samples, the processed RNA-seq data (BAM files) were downloaded from ENCODE and listed in Table S4. The analysis included data from 20 cancer cell lines and 20 tissue samples. For siRNA knock down and SRSF2 mutation samples, the raw data were downloaded from GEO (GSE65349 and GSE164666) and mapped to hg38 using STAR. The resulting BAM files were used for downstream analysis. SGSeq (version 1.36.0) was utilized to identify and quantify splice variants. The splice graph analysis was based on de novo prediction.

**Cell proliferation assay**

At day 0 in the afternoon, cells were treated with trypsin (Sigma, T4090) for digestion. Subsequently, 10 µL of cell suspension was mixed with one part trypan blue (Sigma, T8154) and counted using a cell counter (Bio-Rad, Tc20). Following counting, the cells were seeded onto poly-lysine-coated (Sigma, P4707) 12-well plates at a concentration of $10^4$ cells per well. After reaching the desired time point, the medium was aspirated, and the cells were fixed with 10% formalin (SF100-4) for 5 minutes after being washed with 1 mL of PBS. Subsequently, the cells were incubated with 0.5 mL of 0.1% crystal violet (Sigma, C0775) in 20% methanol for 15 minutes. Following the incubation, the cells were washed three times with PBS. Finally, 1 mL of 10% acetic acid was added to each well to dissolve the crystal violet. Subsequently, 100 µL of the dissolved solution was transferred to a 96-well plate, and the absorbance at 590 nm was measured using a BioTek Synergy 4 spectrophotometer.

**Apoptosis assay**

After trypsinization (Sigma, T4090), 10 µL of cell suspension was mixed with one part trypan blue (Sigma, T8154) and counted using a cell counter (Bio-Rad, Tc20). Following counting, the cells were seeded onto 12-well plates at a concentration of $5\times10^5$ cells per well and cultured overnight. The growth medium was replaced with 1 mL of fresh medium containing 200 ng/mL Fas Ligand (PeproTech, 310-03H) for 4 hours prior to lysis and Western blotting.

**AlphaFold3 Prediction and ChimeraX Visualization**

The interaction between DEDD2 and SLC25A5 was predicted using AlphaFold3 (https://alphafoldserver.com/). Protein sequences were submitted to the server, and the predicted complex structure was downloaded. ChimeraX 1.9 was used for visualization, with DEDD2 and SLC25A5 colored differently to distinguish the proteins.

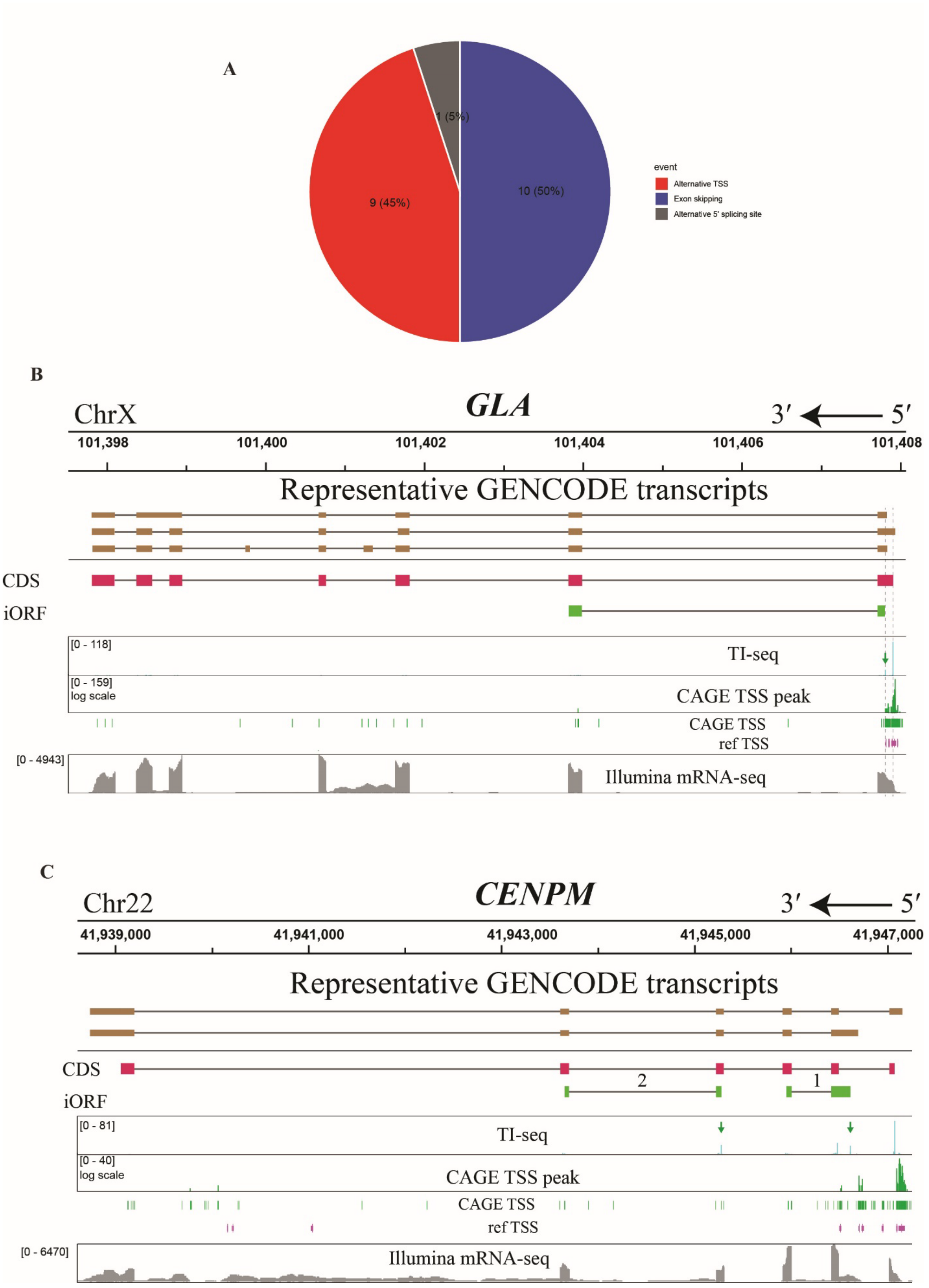
A
1 (5%)
10 (50%)
9 (45%)
event
Alternative TSS
Exon skipping
Alternative 5' splicing site
B
ChrX
GLA
3′ ← 5′
101,398
101,400
101,402
101,404
101,406
101,408
Representative GENCODE transcripts
CDS
iORF
[0 - 118]
TI-seq
[0 - 159]
log scale
CAGE TSS peak
CAGE TSS
ref TSS
[0 - 4943]
Illumina mRNA-seq
C
Chr22
CENPM
3′ ← 5′
41,939,000
41,941,000
41,943,000
41,945,000
41,947,000
Representative GENCODE transcripts
CDS
2
1
iORF
[0 - 81]
TI-seq
[0 - 40]
log scale
CAGE TSS peak
CAGE TSS
ref TSS
[0 - 6470]
Illumina mRNA-seq

**Fig. S1. Alternative transcription start sites (TSS) and alternative pre-mRNA splicing generate iORF-encoding alternative transcripts.** (**A**) Classes of iORF-encoding transcript generation mechanisms for 20 representative genes with evidence in GENCODE for alternative transcripts lacking the full-length CDS. (**B**) In the *GLA* gene, an internal TSS within the first exon bypasses the canonical start codon. Tracks represent translation initiation site sequencing (TI-seq, cyan), CAGE TSS (green), and mRNA-seq (gray) supporting iORF expression. (**C**) In the *CENPM* gene, an alternative TSS generates a transcript initiating at a 5' extended exon 2 that bypasses the CDS start codon. Two iORFs are supported by TI-seq (cyan and green arrows), CAGE-seq (green), and mRNA-seq (gray).

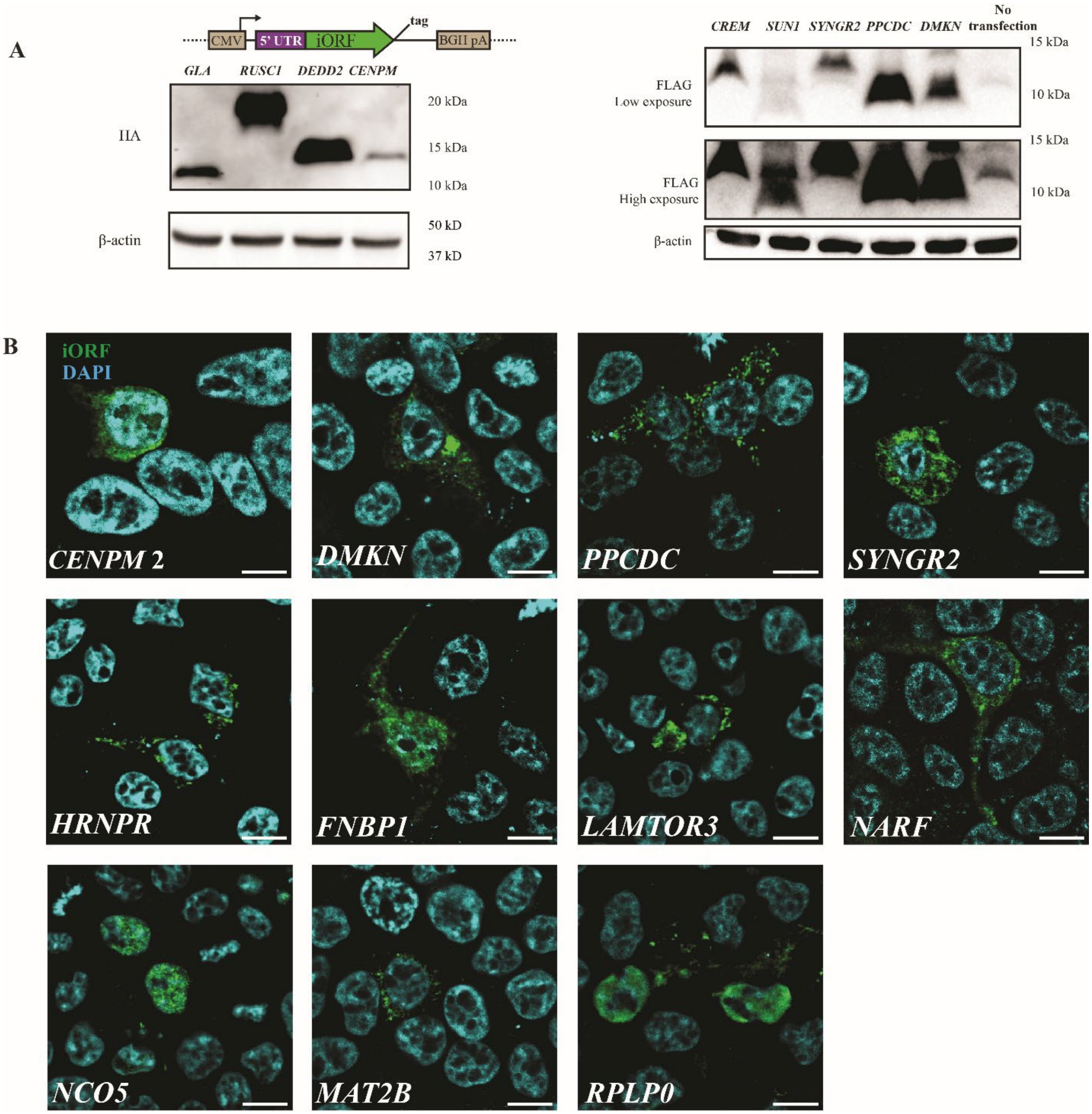


**Fig. S2. iORF translation in overexpression assays.** (**A**) iORF-encoding transcripts (5'UTR to iORF stop codon) were cloned from cDNA libraries into pcDNA3.1 with HA-FLAG tags and transfected into HEK293T cells with empty vector as a control, followed by anti-FLAG Western blot after immunoprecipitation. (**B**) Confocal microscopy of HEK293T cells transfected with epitope-tagged iORF expression plasmids and subjected to anti-FLAG immunofluorescence (green) and DAPI staining (cyan). Scale bars, 10 μm.

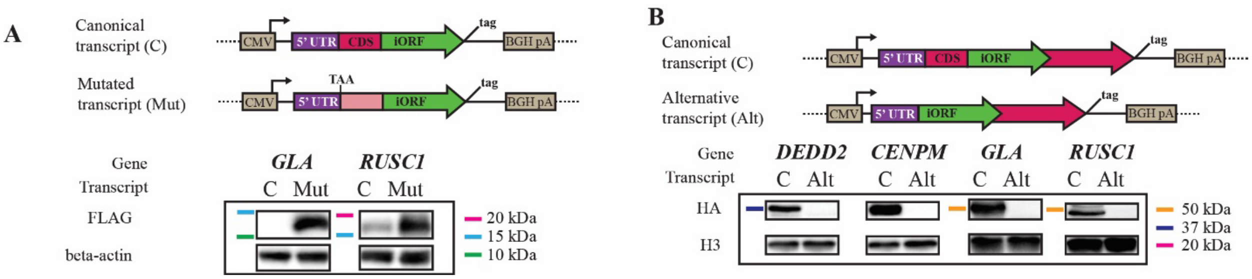


**Fig. S3. Canonical and alternative transcripts uniquely support CDS vs. iORF expression.** (**A**) GENCODE canonical (C) transcripts encoding the CDS from four representative genes were cloned (5' end to iORF stop codon, with FLAG tags appended to the 3' end of the iORF) and transfected into HEK293 cells. CDS start codons were mutated to TAA (Mut) to assess iORF expression in the absence of upstream translational repression present in the canonical (C) transcript. Anti-FLAG Western blot was performed post-transfection to assess iORF translation. (**B**) GENCODE canonical (C) and alternative (Alt) transcripts (5' end to CDS stop codon, with HA tags appended to the 3' end of the CDS) were cloned and transfected into HEK293 cells. Anti-HA Western blotting was performed to assess CDS translation from each context.

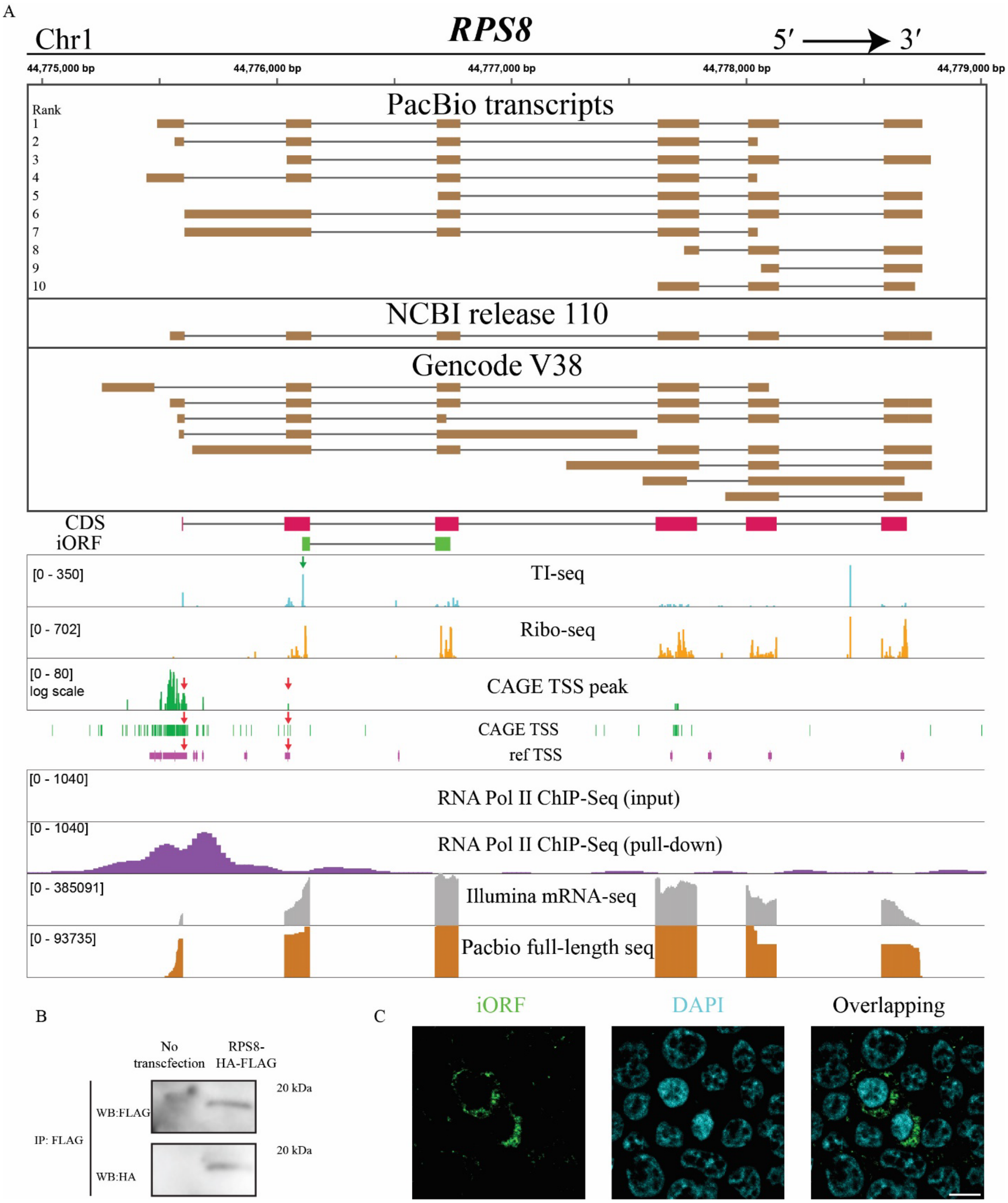


**Fig. S4. Validation of the *RPS8* alternative transcript and iORF translation.** (**A**) Genome view of *RPS8* showing representative transcripts from PacBio sequencing (tracks 1–10, taupe, exons) (highlighting the third-ranked iORF-encoding transcript with a novel 5' end utilizing an alternative TSS), NCBI, and GENCODE. CDS and iORF coding regions are depicted in magenta and green, respectively. TI-seq (cyan) and other data sources (CAGE-seq, reference TSS, PolII ChIP-seq, mRNA-seq, PacBio) support an internal TSS (red arrows) driving alternative transcript expression. (**B**) The top PacBio-detected alternative transcript uniquely encoding the iORF was cloned (5' end to iORF stop codon, fused with dual HA-FLAG tags) and transfected into HEK293T cells. Anti-FLAG Western blotting was performed to assess iORF translation, with β-actin as a loading control and untransfected cells as a negative control. (**C**) Transfected cells from (B) were fixed and stained for anti-FLAG immunofluorescence (green) with DAPI counterstain (blue), showing subcellular localization of the iORF-encoded protein.

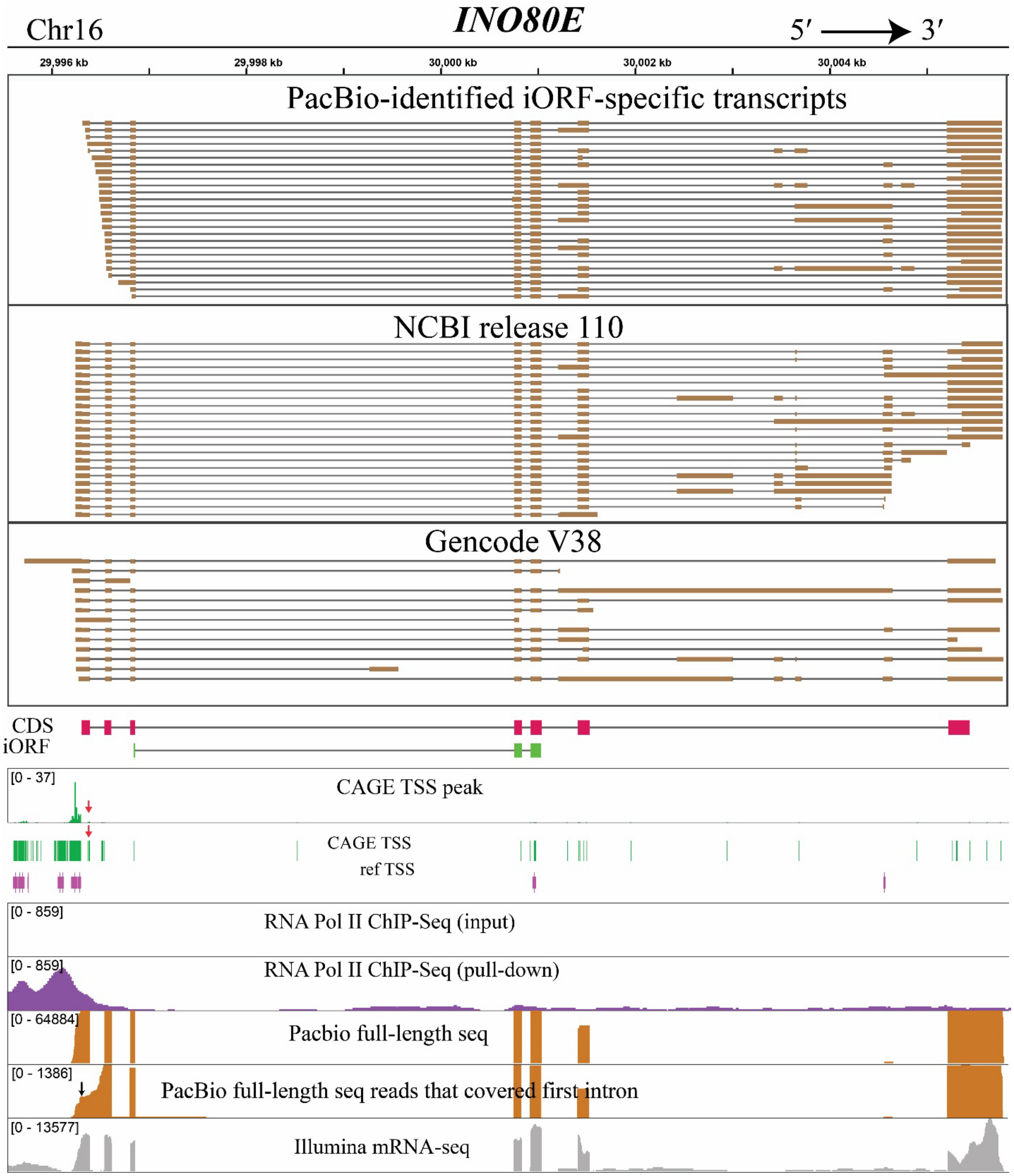


**Fig. S5. PacBio Sequencing identified novel transcript for *INO80E* iORF.** Genome view of *INO80E* showing iORF-specific transcripts from PacBio sequencing (taupe, exons), NCBI, and GENCODE. CDS and iORF coding regions are depicted in magenta and green, respectively. Weak CAGE-seq, reference TSS, diffused PolII ChIP-seq, and PacBio reads support a downstream TSS (red arrows) driving alternative transcript expression. The black arrow indicates the start codon of CDS.

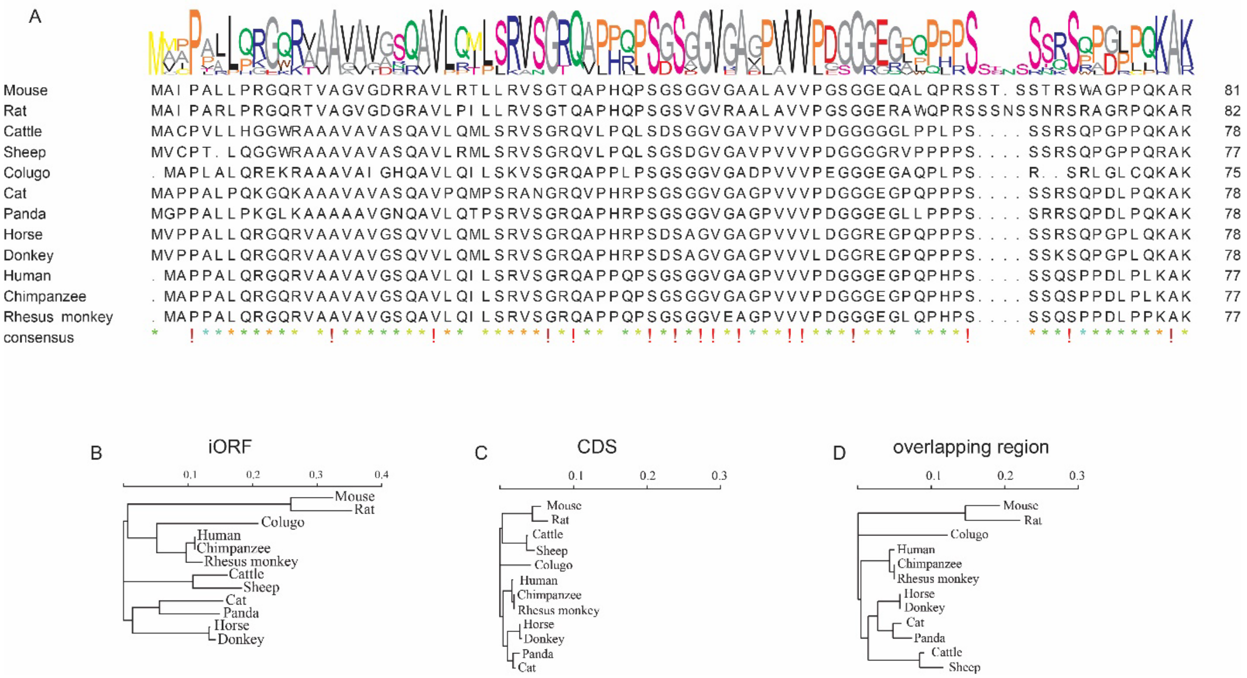


**Fig. S6. Conservation analysis of DEDD2 CDS and iORF.** (**A**) Multiple sequence alignment of DEDD2 iORF sequences across 12 species. (**B**) Phylogenetic tree of DEDD2 iORF sequences, with branch lengths representing evolutionary distances (scale bar: sequence divergence). (**C**) Phylogenetic tree for the DEDD2 CDS. (**D**) Phylogenetic tree for the region of the CDS overlapping the iORF (CDS reading frame), with branch lengths indicating evolutionary distances.

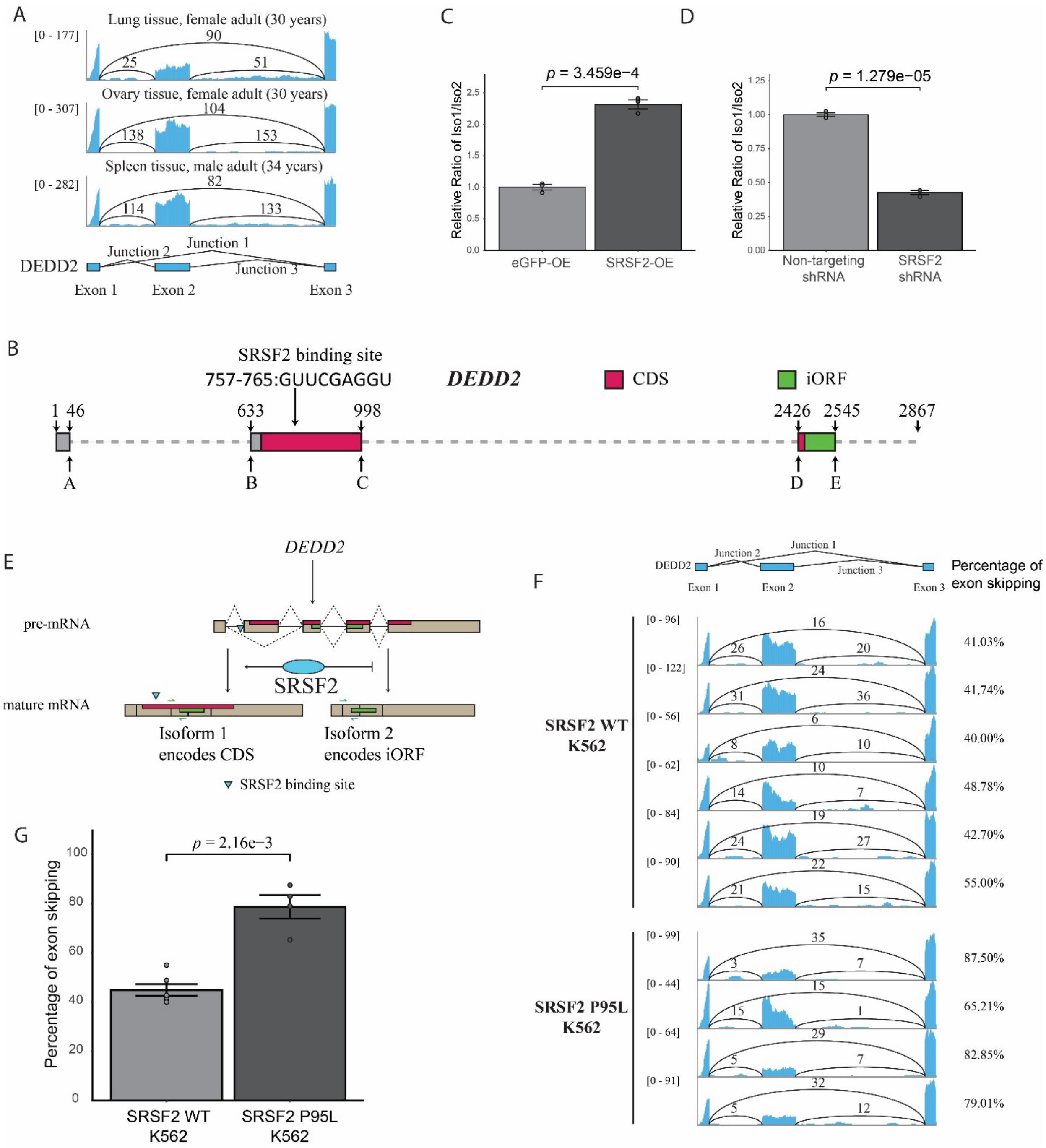


**Fig. S7. SRSF2 regulates DEDD2 splicing.** (**A**) ENCODE RNA-seq read depth and splice junction counts (exons 1–3) for *DEDD2* in normal human tissues. (**B**) Schematic of DEDD2 pre-mRNA (exons 1–3, up to position 2867). Magenta: CDS; green: iORF; arrow: predicted SRSF2 binding site within exon 2. (**C**, **D**) qRT-PCR of DEDD2 transcript variants: exon 2 included (variant 1) vs. excluded (variant 2) under conditions of (C) overexpression of SRSF2 or eGFP control, or (D) shRNA silencing of SRSF2 vs. non-targeting control. ***p***-values by Welch's t-test; bars: mean ± SEM (n = 3). (**E**) Diagram of model for SRSF's role in exon 2 inclusion. (**F**) RNA-seq read depth across DEDD2 exons 1–3 and splice junctions (1–2, 2–3 for variant 1; 1–3 for variant 2) in K562 cells expressing WT or P95L SRSF2. (**G**) Bar plot of exon 2 exclusion percentages in K562 cells with WT or P95L SRSF2. ***p***-values by Welch's t-test; bars: mean ± SEM (n = 6 for WT, n = 4 for P95L).

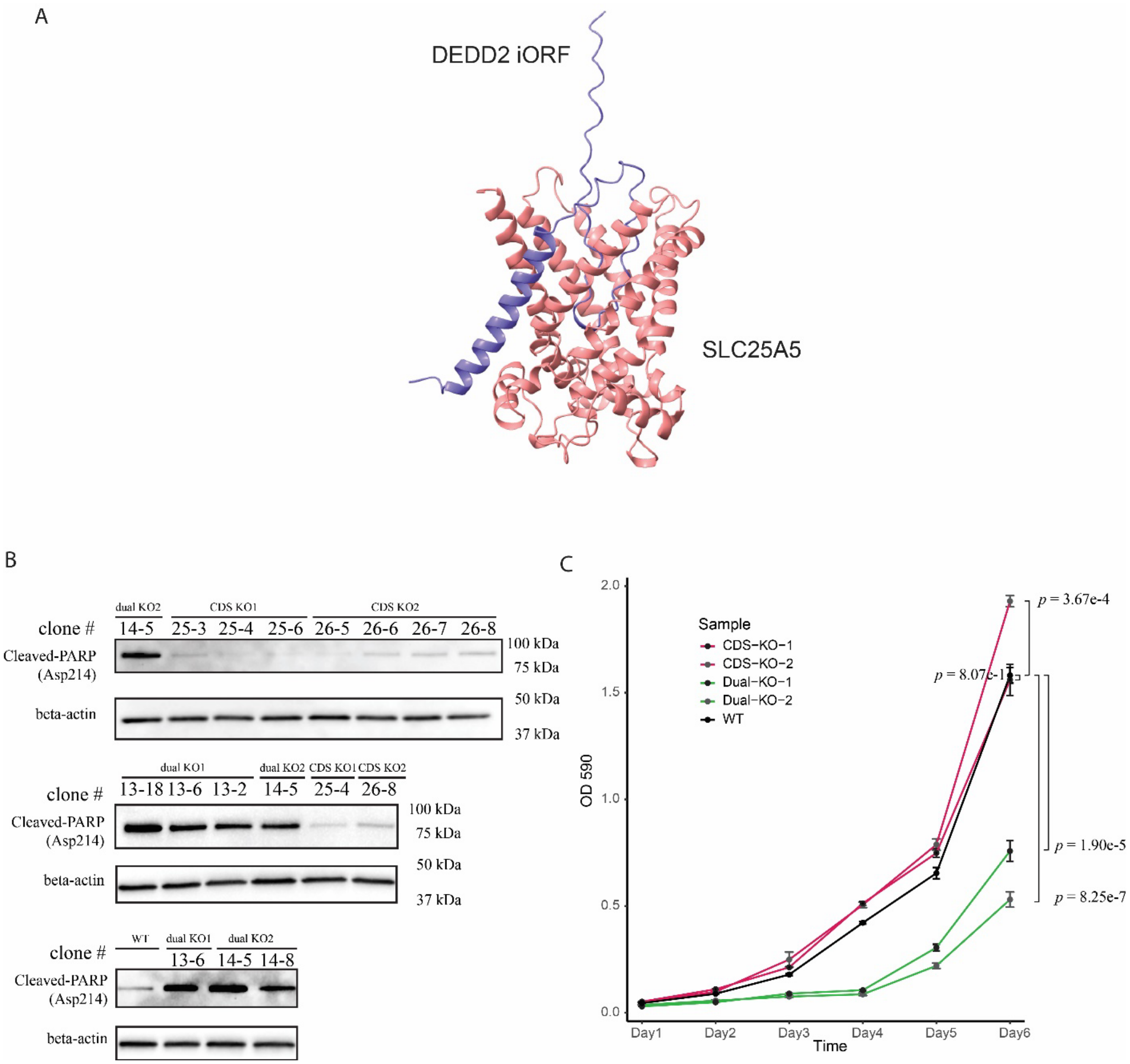


**Fig. S8. DEDD2 validation experiments and controls.** (**A**) AlphaFold 3 predicted interaction between DEDD2 iORF microprotein (purple) and SLC25A5 (pink), with an interface pTM (ipTM) score of 0.22 and a predicted TM-score (pTM) of 0.67. (**B**) Western blot (Fig. 4D) showing cleaved PARP (Asp214) levels in additional clonal CDS KO and dual KO cell lines. Clone numbers are indicated; β-actin was the loading control. (**C**) Cell proliferation assay comparing WT, CDS KO, and dual KO lines (two clones each). Cells were seeded equally and grown for six days; cell numbers were measured via crystal violet staining (OD 590). Error bars indicate mean ± SEM (n = 4). Statistical significance on Day 6 was determined by Welch's t-test.
